# How do gepotidacin and zolifodacin stabilize DNA-cleavage complexes with bacterial type IIA topoisomerases? 2. A Single Moving Metal Mechanism

**DOI:** 10.1101/2024.10.15.618406

**Authors:** Robert A. Nicholls, Harry Morgan, Anna J. Warren, Simon E. Ward, Fei Long, Garib N. Murshudov, Dimitry Sutormin, Benjamin D. Bax

**Affiliations:** Scientific Computing Department, UKRI Science and Technology Facilities Council, Research Complex at Harwell, Rutherford Appleton Laboratory, Didcot, Oxfordshire. OX11 0FA, UK; Medicines Discovery Institute, Cardiff University; Diamond Light Source, Harwell Campus, Didcot, Oxfordshire. OX11 0DE, UK; MRC Laboratory of Molecular Biology, Cambridge; Institute for Systems Biology, Seattle, WA, USA

**Keywords:** Novel Bacterial Topoisomerse Inhibitor (NBTI), spirocyclic pyrimidine trione (SPT), fluoroquinolone, moxifloxacin, gepotidacin, zoliflodacin, type IIA topoisomerase, DNA-cleavage

## Abstract

DNA gyrase is a type IIA topoisomerase that can create temporary double-stranded DNA breaks to regulate DNA topology and an archetypical target of antibiotics. The widely used quinolone class of drugs use a water-metal ion bridge in interacting with the GyrA subunit of DNA gyrase. Zoliflodacin sits in the same pocket as quinolones but interacts with the GyrB subunit and also stabilizes lethal double-stranded DNA-breaks. Gepotidacin had been observed to sit on the twofold axis of the complex, midway between the two four base-pair separated DNA-cleavage sites and has been observed to stabilize singe-stranded DNA-breaks. Here we use information from three crystal structures of complexes of *Staphlococcus aureus* DNA gyrase (one with a precursor of gepotidacin and one with the progenitor of zoliflodacin) to propose a simple single moving metal-ion catalyzed DNA-cleavage mechanism. Our model explains why the catalytic tyrosine is in the tyrosinate (negatively charged) form for DNA-cleavage. Movement of a single catalytic metal-ion (Mg^2+^ or Mn^2+^) guides water mediated protonation and cleavage of the scissile phosphate which is then accepted by the catalytic tyrosinate. Type IIA topoisomerases need to be able to rapidly cut the DNA when it becomes positively supercoiled (in front of replication forks and transcription bubbles) and we propose that the original purpose of the small Greek Key domain, common to all type IIA topoisomerases, was to allow access of the catalytic metal to the DNA-cleavage site. Although the proposed mechanism is consistent with published data, it is not proven and other mechanisms have been proposed. Finally how such mechanisms can be experimentally distinguished is considered.

## 1. Introduction

One of the aims of modern structural biology is to understand how movement is coupled to the chemical making and breaking of bonds, that is to “watch chemistry happen” [1]. In physics, ‘the three body problem’ pertains to describing the motion of three bodies under gravitational attraction [2]. ‘ *At first glance, the difficulty of the problem is not obvious, especially when considering that the two-body problem has well-known closed form solutions given in terms of elementary functions. Adding one extra body makes the problem too complicated to obtain similar types of solutions. [*…*] solutions do not exist because motions of the three bodies are in general unpredictable, which makes the three-body problem one of the most challenging problems in the history of science [2]*.’ There is an analogous problem in chemistry, where the Schrödinger equation can be solved analytically for the hydrogen atom, owing to the simplicity of the two-particle physical system [3]. This explains why physics has so far failed to properly explain catalytic chemistry, and why experimental structural science is still required.

One of the long-standing challenges for structural biology is a catalytic mechanism of topoisomerases – complex and flexible enzymes which modify the topology of DNA. Topoisomerases are divided into two classes; type I and type II (see supplementary Figure S1). While type I topoisomerases use single-stranded DNA breaks to modify DNA topology, type II use double-stranded DNA breaks [4-6]. The introduction of double-stranded DNA breaks is potentially lethal to cells and is normally carefully controlled. Type IIA topoisomerases (Figure 1) include two human enzymes, topoisomerase IIα and IIβ, as well as the bacterial enzymes topoisomerase IV and DNA gyrase. Interestingly, *Mycobacterium tuberculosis* only has DNA gyrase (i.e. it lacks topoisomerase IV) and a type IA topoisomerase [7, 8], which is mechanistically similar. Figure 1, panel b, shows a simplified view of the function of a type IIA topoisomerase. A double-stranded break is made in one segment (the gate or G-DNA duplex) and another DNA-duplex is passed through that break. This is accompanied by large movements of the enzyme [4, 9, 10].

**Figure 1.**
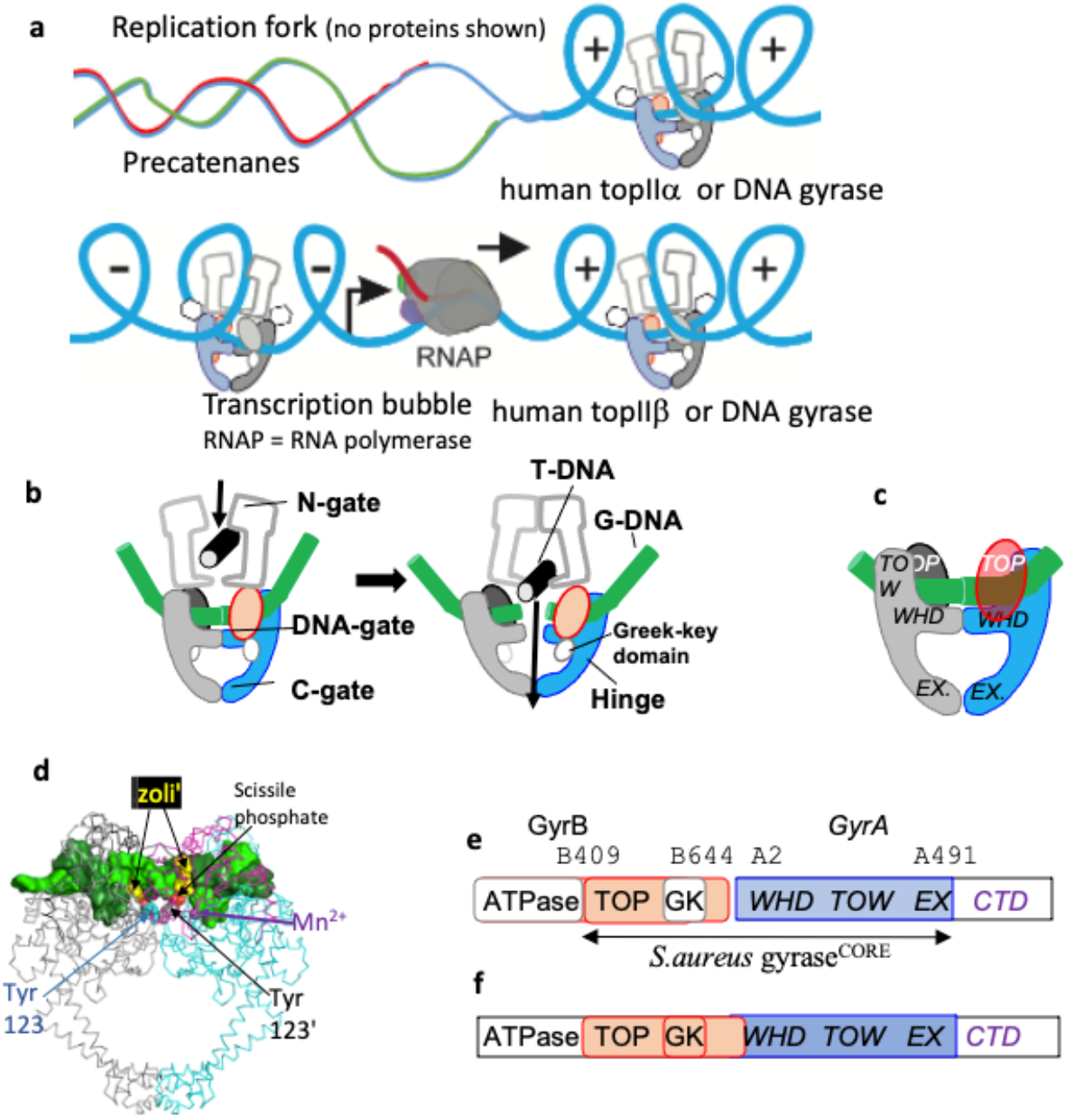
*S. aureus* DNA gyrase is a type IIA topoisomerase. **(a)** Type IIA topoisomerases work ahead of replication forks (and transcription bubbles, at least for long genes [20]) to remove positive DNA-supercoils [21-24] to allow both DNA-replication (and DNA-transcription) to take place. **(b)** Simplified schematic of ATP-dependent relaxation of negatively supercoiled DNA carried out by a type IIA topoisomerase. The gate or G-DNA (green cylinder) is cleaved and another DNA duplex, the T (or transport segment - black) is passed through the cleaved DNA before religation. **(c)** A schematic of *S. aureus* DNA gyrase GyrB27–A56(GKdel) DNA complex with uncleaved DNA (green). One fusion truncate is shown in red and blue, the other in grey and black. Note the small Greek Key domain (residues 544–579) in the GyrB subunit has been deleted and replaced with two amino-acids. **(d)** View of a 2.8Å crystal structure of a GyrB27–A56(GKdel) complex with zoliflodacin (zoli’) and doubly cleaved DNA (PDB code: 8bp2). The protein is shown as a Cα trace with GyrA subunits cyan or grey and GyrB subunits magenta or black; the cleaved DNA (surface representation) is colored green or forest green (a similar coloring scheme is used throughout this paper). The side-chains of the catalytic tyrosines, which are covalently bonded to the cleaved DNA via the scissile phosphates, are shown in sphere representation GyrA Tyr 123 (cyan) and Tyr 123’ (grey). Zoli-flodacins are shown as spheres (yellow). An Mn^2+^ ion at the Y (or B) site is shown as a purple sphere. **(e)** DNA gyrase consists of two subunits, GyrB and GyrA. The *S. aureus* DNA gyrase GyrB27– A56(GKdel) construct used to determine many crystal structures is a fusion of the C-terminal Toprim (TOP) domain of GyrB with the winged helical domain (WHD), tower (TOW) and exit-gate (EX) domains from GyrA. The C-terminal domain (CTD), which wraps DNA in DNA gyrase, is not shown in panel b (the CTD has different functions in other type IIA topoisomerases). **(f)** In humans (and yeast) topoII is a single subunit and functions as a homodimer.

How the two bacterial type IIA topoisomerases, DNA gyrase and topoisomerase IV, cleave DNA and modulate DNA topology is of interest because three classes of antibiotics stabilizing DNA-cleavage complexes with these targets have now successfully passed phase III clinical trials. The flexibility these enzymes require, in making a double-strand DNA-break in one DNA-duplex and then passing a second DNA-duplex through that break, has caused problems in obtaining high resolution crystal structures. A clear definition of how the catalytic metal ion(s) regulate this DNA-cleavage process in a normally safe manner follows from high-resolution (better than 2.2Å) crystal structures [11]. The deletion of the small Greek Key domain in *S. aureus* DNA gyrase improved the resolution of crystal structures with GSK299423 (a precursor of gepotidacin) from 3.5 Å (with the Greek Key domain) to 2.1 Å (with the Greek Key domain deleted) and gave a clear view of a catalytic metal at the 3’(A) position [12]. This *S. aureus* DNA gyrase Greek Key deletion construct has been successfully used in a large number of crystal structures [11, 13], but has not yet been successfully replicated in other type IIA topoisomerases - which all contain the small Greek Key domain.

An initial structure of moxifloxacin with *A. baumanii* Topoisomerase IV in a DNA-cleavage complex [14] showed the presence of the water-metal ion bridge [15] and explained two common target-mediated resistance mutations. However, the DNA used in this initial structure was from a previous lower-resolution structure [16] and the limited resolution of the data (3.25 Å) precluded a clear visualisation of the register of the DNA [14]. Subsequent higher-resolution structures with a shorter version of this DNA, quinolones and a *Mycobacterium tuberculosis* DNA gyrase construct [17] clearly showed that the asymmetric DNA sequence was in two orientations in the crystal - related by the twofold axis of the complex. The register of the DNA sequence in the initial *A. baumanii* Topoiso-merase IV structure is uncertain [14]. A 2.95Å structure of the *S. aureus* DNA gyrase Greek Key deletion construct (Supplementary Figures S2 and S3) in a DNA-cleavage complex with moxifloxacin (pdb code: 5cdq [18]), used a 20 base-pair symmetric (palindromic) DNA sequence to avoid the possibility of the DNA suffering from static disorder around the twofold axis of the complex [19]. This moxifloxacin structure (5cdq-BA-x.pdb - Supplementary Figure S3) is very similar to the moxifloxacin DNA-cleavage complexes with *A. baumannii* Topo IV [14]. However, the 2.95 Å structure is with the Greek Key deletion mutant (see Supplementary Figure S2). The water-metal ion bridge [15] is conserved in moxifloxacin DNA-cleavage complex structures between *A. baumanii* Topoisomerase IV [14] and *S. aureus* DNA gyrase [18] and in a modified form in *M. tuberculosis* DNA gyrase [17], (see Figure 3 in [17]-note the presence of an alanine at position 90 in *M. tuberculosis* GyrA cf serine in other bacteria). Interestingly the *Mycobacterium tuberculosis ‘genome encodes only one copy of type I and one copy of type II topoisomerase’* [7].

An extensive corpus of biochemical and structural information strongly suggests that type IIA topoisomerases require a catalytic metal (preferentially Mg^2+^) in the Toprim domain for proper DNA cleavage and religation; while type IA topoisomerases seem to only require the metal ion for religation by the Toprim domain [8], DNA-cleavage being accomplished with the aid of a lysine residue. Type IA topoisomerases only work on negatively supercoiled DNA [8]. There are currently three theories (see Table 4 and discussion in [11]) of how the G-segment DNA is cleaved by type IIA topoisomerases such as DNA gyrase: (i) a two metal mechanism which was proposed [25] based on a misinterpretated electron density map [25]. To date, to the best of our knowledge, there has been no structural data published that is consistent with such a model, and therefore it is unlikely to be correct (ii) a single-metal mechanism in which DNA-cleavage is proposed to take place when the metal ion is at the 3’(A) site described (pages 3438-3447) in [13]. This model is also unlikely to be correct [26, 27]; noting that it does not explain why the catalytic tyrosine is in the tyrosinate (negatively charged) form, nor explain how the enzyme uses supercoiling to control DNA-cleavage and (iii) the single moving metal mechanism described in this paper, which explains why the catalytic tyrosine is in a tyrosinate (negatively charged) form for DNA-cleavage, and which also proposes that the small Greek Key domain controls metal access to the catalytic site to allow rapid relaxation of positively supercoiled DNA ahead of replication forks and transcription bubbles (see Figure 1). In this paper a chemically sensible single moving metal mechanism [26, 27] is proposed, based on three published crystal structures [12, 18, 28]. The aim of this paper is not to prove a mechanism, but to present a new mechanism consistent with existing data (including chemistry). Experiments are then proposed to “watch chemistry happen”.

Metal binding Toprim domains were named for topoisomerases and primases and were characterized in a multiple sequence alignment in a 1998 paper [29]. However, the first Toprim domain in a type IIA topoisomerase was described in the structure of the catalytic core of yeast topoisomerase II published in a paper in 1996 [30]. This stated that in addition to the Toprim domain there is a simple Greek key domain ‘*(residues 565-605) is inserted between the fifth strand and final helix* [of the Toprim domain]’. Although Toprim domains are found in many other proteins, including other topoisomerases (e.g. type IA topoisomerases) the insertion of the simple Greek Key domain between the fifth strand and subsequent helix of the Toprim domain seems to be a feature which is found uniquely in all type IIA topoisomerases. In Gram negative DNA gyrases (such as *E. coli* DNA gyrase), the Greek Key domain is augmented by an inserted domain of some 174 residues; this inserted domain is not present in other type IIA topoisomerases, which have only the small Greek Key domain inserted in the Toprim domain (see sequence alignment in Supplementary Figure S4). The small Greek Key domain in type IIA topoisomerases tends to be mobile - it was not seen in the 2.15Å structure of etoposide with human topo IIB [31].

A *S. aureus* DNA gyrase fusion construct (Figure 1) in which the small Greek Key domain was deleted (residues 544-579 replaced with two amino acids) [12], allowed a clear visualization of a 3’(A) site metal (with a YtoF mutant and GSK299423). A long-standing question has been why type IIA topoisomerases have the additional Greek Key domain within the Toprim domain. One of the conclusions of this paper is that the Greek key domains within type IIA topoisomerases first arose to allow control of metal access to the active site - specifically allowing rapid access of metal ions to control relaxation of positively supercoiled DNA in front of replication forks [21-24].

### 2. Results

In developing the mechanism/movie proposed in this paper we use just four complexes from three crystal structures (see Supplementary Figure S5): (i) the 2.1Å ternary complex with GSK299423 (a precursor of gepotidacin) and uncleaved DNA (PDB code 2xcs, [12]), which has the catalytic tyrosine mutated to phenylalanine and gives a clear view of a 3’(A)-site metal ion (see Table 1); (ii) A 2.5Å DNA-cleavage complex stabilized by QPT-1 (PDB code 5cdm [18]), the progenitor of zoliflodacin (Figure 2 panel c), which has a single metal ion observed at the Y(B) site and (iii) the two complexes (<u>6fqv-c1.pdb</u> and <u>6fqv-c2.pdb</u>) from the 2.6Å binary complex (PDB code 6fqv). This 2.6Å binary structure has two complexes in the asymmetric unit (<u>6fqv-c1.pdb</u> and <u>6fqv-c2.pdb</u> are both available from the ‘Research’ tab at https://profiles.cardiff.ac.uk/staff/baxb); the complexes are similar apart from the position of the catalytic tyrosine (see Figure 2). Neither complex has a metal ion seen at an active site [28].

**Table 1.**
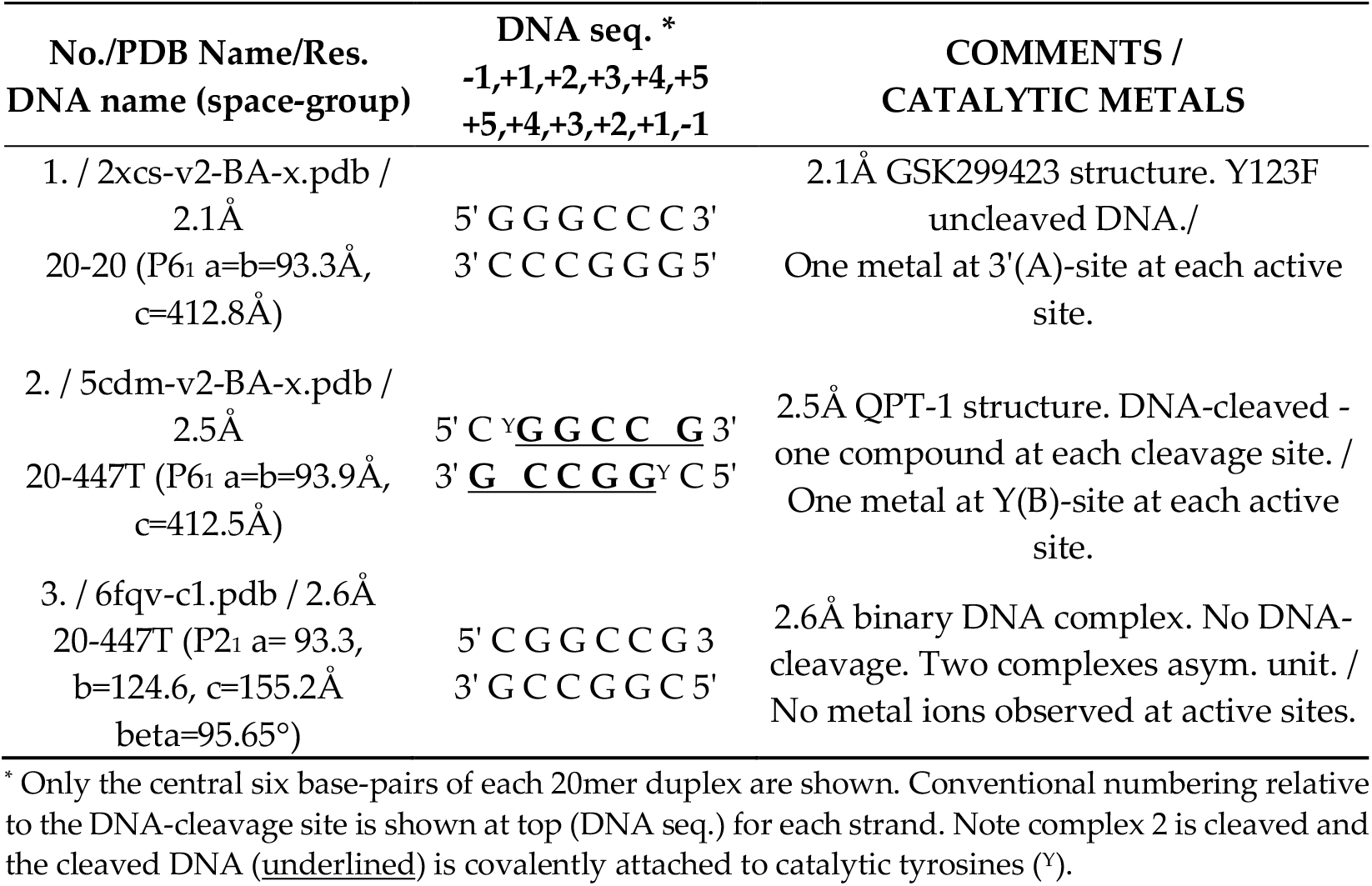
Three crystal structures used in this paper (see [11] for details of (re-)refinement of the two metal ion containing P6_1_ crystal structures).

**Figure 2:**
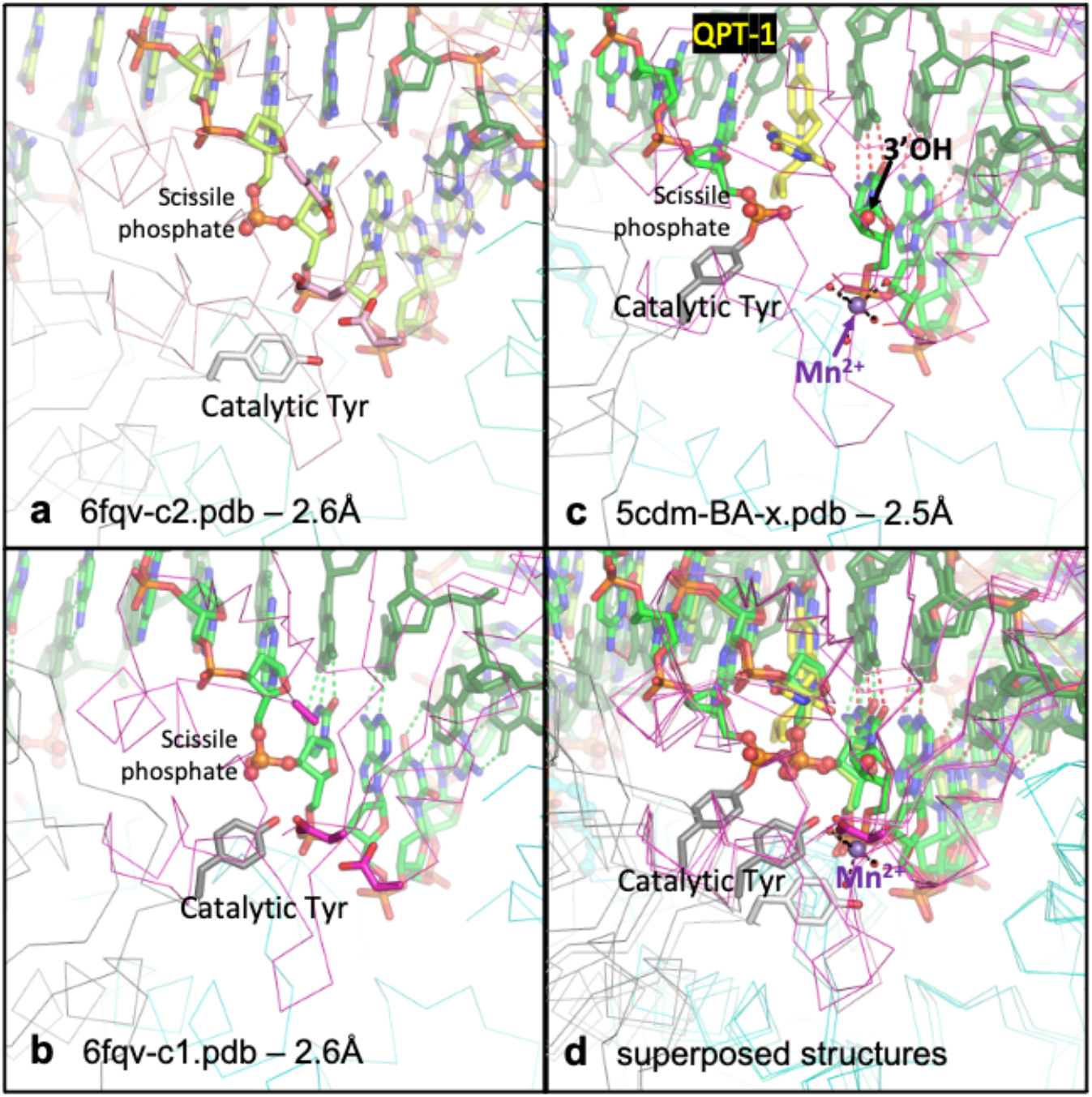
The catalytic tyrosine appears to move. **(a)** View of binary complex 2 (6fqv-c2.pdb. Note that the conformation at the second active site of this - and other - complexes is similar to that at the first active site). **(b)** View of binary complex 1 (6fqv-c1.pdb). **(c)** Equivalent view of 2.5Å QPT-1 complex (5cdm-BA-x.pdb). Note in this structure the catalytic tyrosine has cleaved the scissile phosphate and the QPT-1 sterically inhibits the approach of the scissile phosphate to the 3’OH for religation of the DNA. The metal ion (Mn^2+^) is at the Y (B) site. **(d)** The three structures, from panels a, b and c are shown superposed. Note the relative positions of the catalytic tyrosine – suggestive of its movement. [Structural figures in this paper were created using PYMOL [33].]

**Figure 3.**
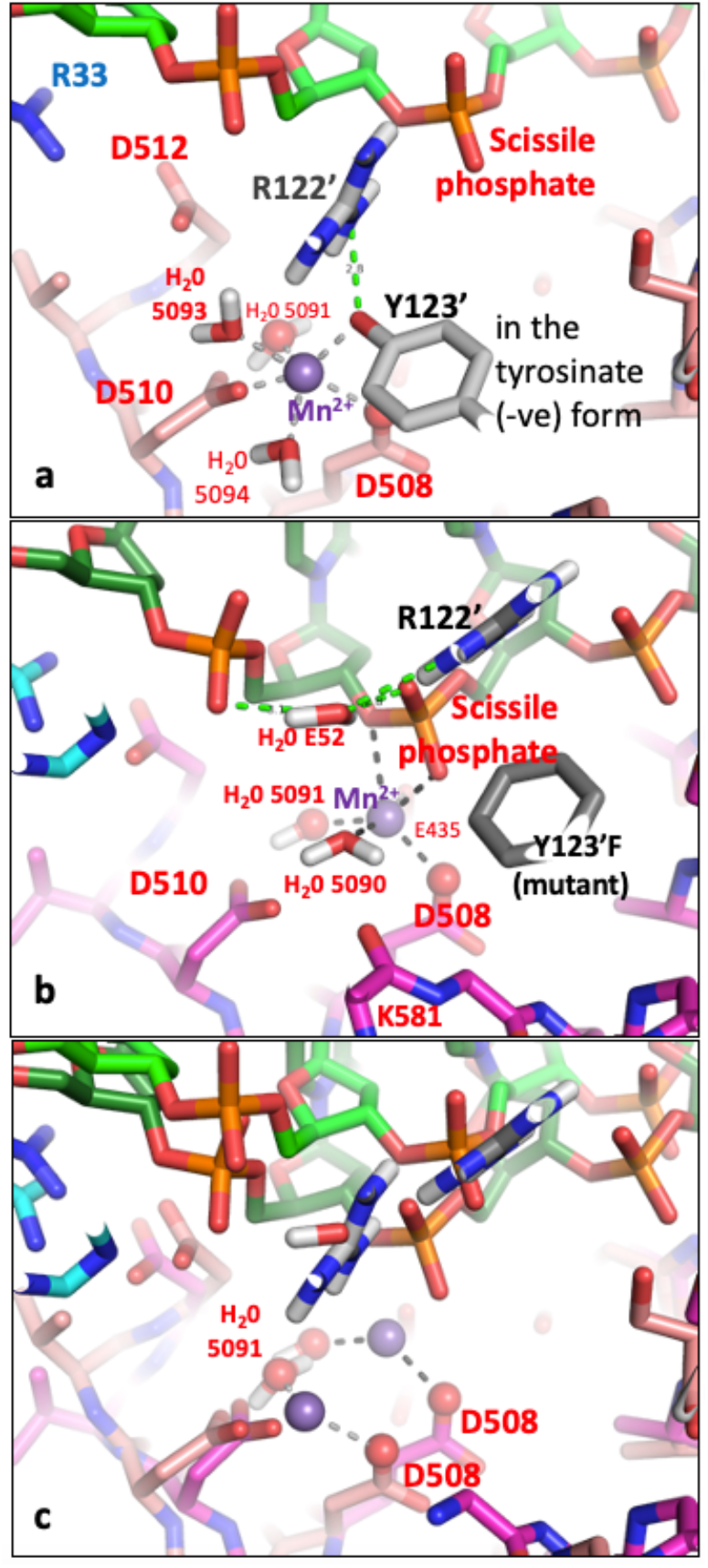
A comparison of state2.pdb with 2xcs-v2-two-DNA.pdb – only some hydrogens (white atoms) are shown. **(a)** Tyrosine 123’ from state2.pdb (derived from 6fqv-no1-one-DNA.pdb) is shown accepting a hydrogen bond from arginine 122’ and also coordinating a catalytic Mn^2+^ ion (at the Y(B) site - the terminal oxygen of the tyrosine is at a similar position to water 5095 in 5cdm – see also Supplementary Figures S7 and S8). The catalytic tyrosine is in the tyrosinate form with a negative charge on the terminal oxygen. **(b)** In the 2.1Å 2xcs-v2-two-DNA.pdb a single Mn^2+^ ion is observed coordinated by two waters, Asp 508 and Glu 435 as well as two oxygens from the scissile phosphate. (see Figure 5 in [12]). Water E52 donates hydrogen bonds to the scissile phosphate and the previous phosphate and accepts a hydrogen bond from the NE of Arg 122’. This E52 water is a new feature of the binding pocket - it is some 5.5Å from Mn^2+^ 5081 (the equivalent water, F41, is some 5.3Å from Mn^2+^ D5081). Only some hydrogens (white atoms) are shown. **(c)** Superposition of structures from (a) and (b). Note the Mn^2+^ ion moves some 3.2Å between the superposed structures; it remains coordinated by the side chain of Asp 508 (which moves 2.2 Å) and the ‘inside water’ H_2_O 5091 (moves 1.1 Å). Other waters (except E52) and Y123’ (Y123’F) and K581 have been removed from the picture for clarity. Note Arg 122’ also moves – its position in (a) is similar position to E52 from (b).. Only preserved coordination sphere lines from OD2 Asp 508 and oxygen of water 5091 are shown to Mn^2+^ ion (purple sphere).

Before a movie could be made the crystallographic coordinates (see Table 1) had to be adjusted so that: (i) The structural models had the same DNA sequences (in each of four sets of coordinates used in making the movie); (ii) Each set of coordinates contained a single molecule (electron density maps typically represent thousands of molecules in the crystal, so side-chains where there was electron density for more than one conformation, or no visible supporting electron density, were modelled as single conformations); (iii) Hydrogen atoms were added to structures using the program Maestro [32] (see Materials and Methods for details).

#### 2.1 The catalytic tyrosine seems to move

To model chemistry happening - based on multiple crystal structures (Table 1), it is useful if the catalytic metal ion has the same name in different crystal structures. To do this the standard BA-x nomenclature is used for *S. aureus* gyrase PDB files (see Materials and Methods and [11] for details of this naming system). The catalytic tyrosine is residue 123 in *S. aureus* DNA gyrase = 122 in *E.coli* DNA gyrase - (see supplementary Figure S6).

In Figure 2 the position of the catalytic tyrosine is shown in three different positions, in three different complexes. In 6fqv-c2.pdb (Figure 2a) the catalytic tyrosine (from the second complex in the asymmetric unit from 6fqv.pdb) is not in the catalytic pocket. Whereas in 6fqv-c1.pdb (Figure 2b), that catalytic tyrosine has a position where it can displace a water from the normal coordination sphere of a Y(B)-site metal. However, no metal ions are observed in the catalytic pocket in either complex in the 6fqv crystal structure [28]. This seems to be because the phosphate prior to the scissile phosphate is too far away from the metal binding site to interact with a water. In high-resolution crystal structures where a Y(B)-site metal is observed (e.g. [31]), a water molecule is observed sitting between the catalytic metal ion and the phosphate prior to the scissile phosphate (from the DNA). In this paper the Y(B), 3’(A) nomenclature is used for the two observed metal positions (see [11] for details). This is because, as shown in Figure 2, the catalytic tyrosine appears to move.

When the catalytic tyrosine cleaves the DNA (as shown in Figure 2c) it becomes covalently attached to the scissile phosphate. Note that the active sites are formed in trans with the catalytic tyrosine C123’ from the C subunit (of the DC-fusion protein) cleaving the DNA with the B-subunit (from the BA-fusion protein) Toprim domain (and, vice versa, tyrosine A123 cleaving with the D-subunit Toprim domain). Standard crystallographic nomenclature where residues from the first (BA) chain are not given a subscript, whereas residues from the second (DC) chain are (e.g. C123’ or just 123’) is used below in this paper.

#### 2.2 A moving metal mechanism for DNA-cleavage

The position of the catalytic tyrosine in 6fqv-v1.pdb is such that its terminal oxygen is within 1.6 Å of water 5095 in 5cdm (see Figure 2 and Supplementary Figure S7); i.e. the terminal oxygen of the tyrosine and water 5095 occupy a similar position in space. Because the catalytic tyrosine appears to move (see Figure 2) the terminal oxygen of the tyrosine can be moved about 1.6 Å back towards its position in 6fqv-c2.pdb (Figure 2) to be closer to the position of water 5095 - and displace it from the Mn^2+^ coordination sphere (see Materials and Methods; Tables 2 and 3). In this starting position the tyrosine is accepting a hydrogen bond from Arg 122’, but the slight move backwards means Arg 122’ is too far from the scissile phosphate to donate a hydrogen bond to the scissile phosphate. The catalytic tyrosine (A123 or C123’) is believed to be in the tyrosinate (O-rather than OH) form before it cleaves the DNA (see discussion). In cleaving the DNA the tyrosinate form is stabilized by interactions with both the catalytic metal (Mn^2+^) and the positively charged side chain of Arg122 (Figure 3a). [Arg A122 (or Arg C122’) is an important catalytic residue but is often quite mobile in *S. aureus* gyrase crystal structures].

**Table 2.**
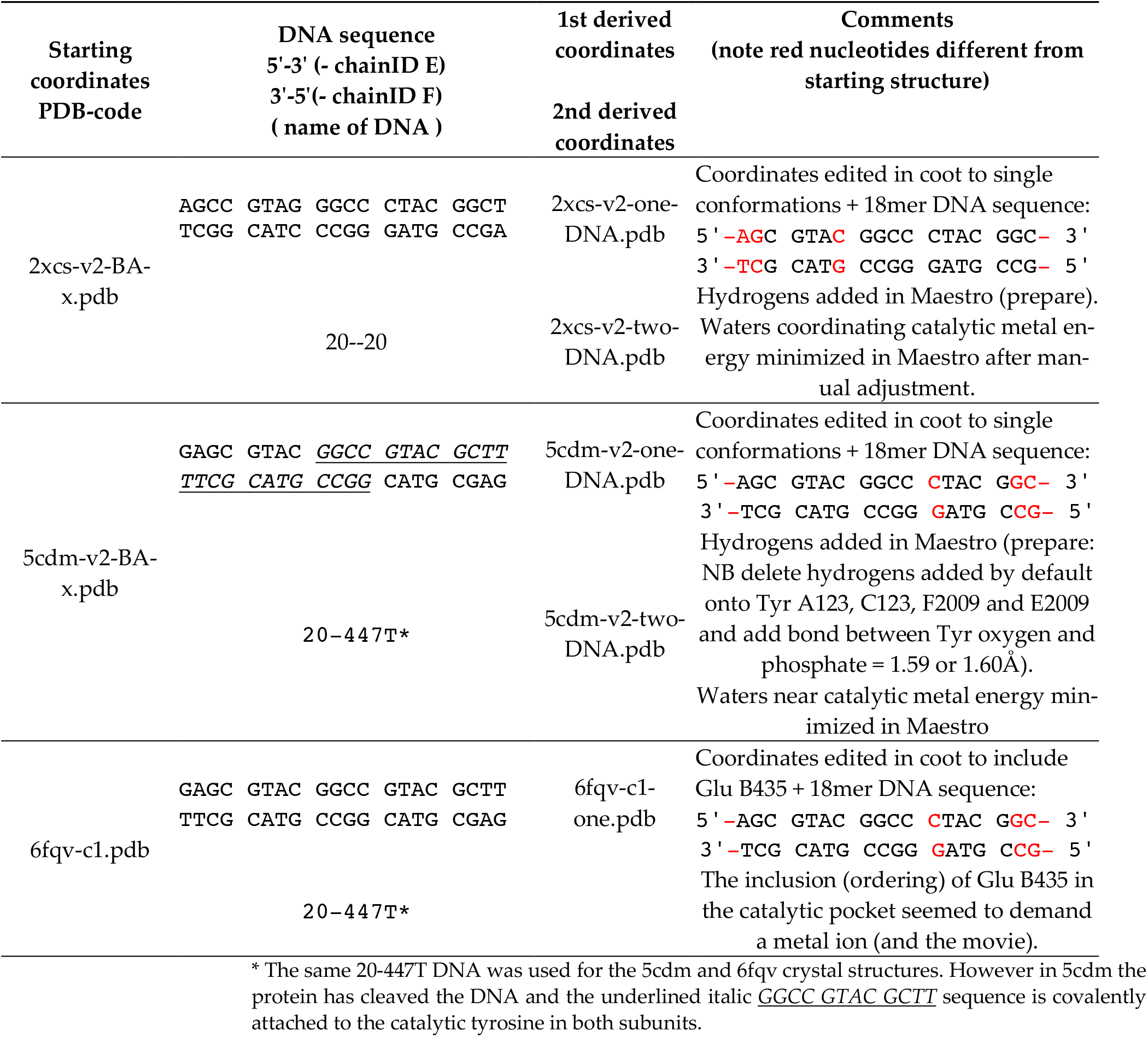
Derived coordinates in preperation for construction of a molecular movie.

**Table 3.** Coordinates used in making the movie.

| No./PDB Name | COMMENTS |
| --- | --- |
| 1. state1.pdb | Derived from 6fqv-c2.pdb. Contains no catalytic metal ion (see Figure 2a). |
| 2. state2.pdb | Derived from 6fqv-c1-one.pdb. Metal ion 5081 and waters 5091, 5093 and 5094 added. The catalytic tyrosine moved slightly so terminal oxygen sits where water 5095 sits in 5cdm (see Figure 3). |
| 3. state3.pdb | Derived from **state2.pdb**. Moved towards 2xcs-v2-two-DNA.pdb. Water 5095 is based on water E 52. DNA not yet cleaved. |
| 4. state4.pdb | Derived from **state3.pdb**. DNA-cleaved as in 5cdm-v2-two-DNA.pdb. Water 5095 has come in from position in state 3.pdb. |

The model presumes that the metal comes in as the catalytic tyrosine moves. One possibility is that the metal access to the active site may be controlled by the Greek Key domains for positively supercoiled DNA (see Conclusions).

In the proposed DNA-cleavage mechanism the Y(B) site metal is coordinated by the catalytic tyrosine (Y) prior to DNA-cleavage (Figure 3a) and the metal is then attracted towards the 3’(A) site (Figure 3b) but the bond between the 3’ oxygen and the phosphorous atom of the scissile phosphate is cleaved just before the metal reaches the 3’(A) site (Figure 4). Figure 4b suggests that the hydrogen atom from water 5093 is, when transferred to the 3’ oxygen, between this oxygen and the scissile phosphate. However, we now believe the hydrogen on the 3’ oxygen is attracted towards Glu B435 and a hydrogen bond forms between this 3’OH and Glu B435 on DNA cleavage (see Nicholls-et-al-movie1.mp4).

**Figure 4.**
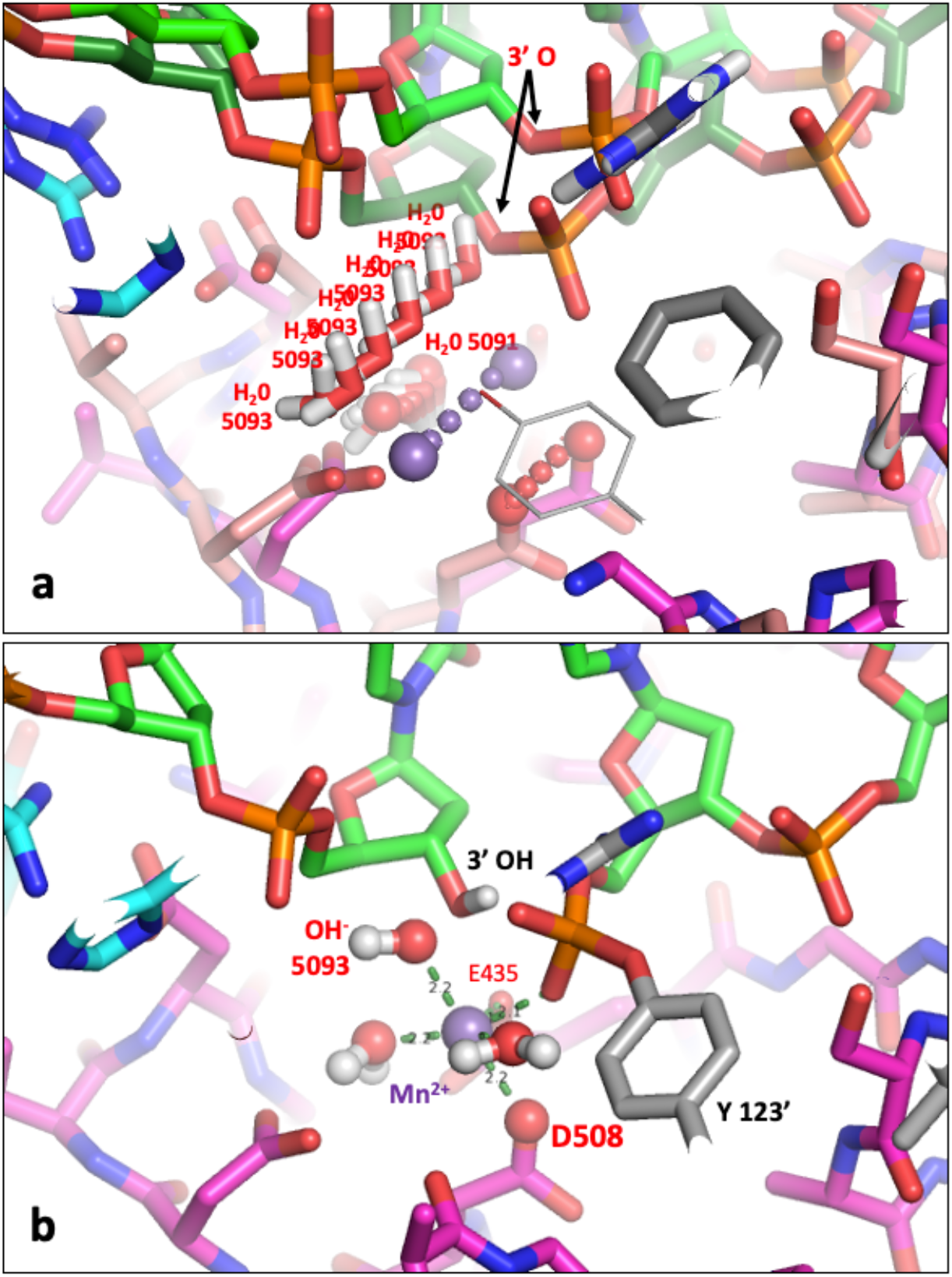
Does water 5093 protonate the 3’-oxygen to cleave the DNA? **(a)** Three conserved atoms, the catalytic metal ion (purple), the OD2 from Asp 508 and the oxygen from water 5091 have been used to define movements and suggest that water 5093 moves to protonate the 3’ oxygen to initially cleave the DNA. Some atoms have been removed from the picture for clarity. **(b)** The Mn^2+^ ion (purple sphere) has moved towards the A-site – thus causing water 5093 to move and protonate the 3’ oxygen from the scissile phosphate, which then becomes the 3’ OH – while water 5093 becomes a hydroxyl ion (OH^-^). In this model the catalytic tyrosine (Y123’) has just accepted the scissile phosphate – while the catalytic metal ion (Mn^2+)^ has an octahedral coordination sphere (two waters, oxygens from Asp B508 and Glu B435 side chains, an oxygen from the scissile phosphate and the oxygen from the OH^-^ 5093 anion). The oxygen atom on D508, which maintains contact with the metal ion throughout the reaction, is shown as a red sphere. Water 5095 is not shown.

Note in the binary complex, 6fqv, the side-chain of Glu 435 is disordered; presumably it tries to get away from the scissile phosphate and the side chains of Asp 508 and Asp 510. In structures containing a metal ion in the active site Glu 435 becomes ordered. In the DNA-cleavage mechanism (Figure 4), on entering the active site the metal ion (Mg^2+^ or Mn^2+^) is attracted towards two negative charges, on the scissile phosphate and on Glu 435, and as it moves it maintains its contact with the side chain of Asp 508 (Figure 3c). Water 5093 which protonates the 3’ oxygen to effect DNA-cleavage, moves as the metal ion moves (Mn^2+^ - purple sphere - Figure 4a).

After the DNA has been cleaved (Figure 4) the DNA gate can be pushed open by a traversing T-DNA duplex, while the metal ion moves back to the Y(B) site, where it acquires a new water - water 5095 - replacing the tyrosine oxygen in the metal ion coordination sphere (see Supplementary Figure S8). Once the T-DNA has passed through the doubly cleaved DNA - presumably the metal at the B-site can religate the DNA.

It is possible, in principle, to propose a simple religation scheme, where water 5095 moves to protonate the tyrosine oxygen, regenerating the tyrosine from the phosphotyrosine. The metaphosphate-like intermediate is presumably then attacked by the 3’ O^-^ ion to regenerate the intact DNA. However, such a religation mechanism has some complications. The 3’OH probably loses a proton to Glu 435, before becoming the attacking 3’ O^-^ ion, but more problematical is the exact conformation of the phosphotyrosine when it is protonated by water 5095. We have not yet fully elucidated such a scheme.

## 3. Discussion

The scheme for DNA-cleavage shown in Figure 4 suggests that the initial step in the cleavage of the DNA is the protonation of the 3’ oxygen. This contrasts with many DNA-cleavage mechanisms which propose that the initial step is caused by a tyrosinate ion attacking a tetrahedral negatively charged phosphate (e.g. Figure 4c in a paper on a type IA topoisomerase [34]). The nucleophilic attack of a negative tyrosinate ion on a negatively charged phosphate ion seems chemically unlikely. The mechanism proposed in this paper for *S. aureus* DNA gyrase suggests that (Figures 4 and 5) - the DNA is first cleaved by protonation of the negatively charged scissile phosphate and then the extremely reactive metaphosphate-like planer intermediate is attacked by the catalytic tyrosinate (Figure 5 panel c). The mechanism proposed in this paper is informed by the non-enzymatic hydrolysis of methylphosphate mechanism (Figure 10.1 and Scheme 10.1 in Frey and Hegeman [35]). The proposed mechanism is also informed by previously proposed mechanisms for phosphotransfer reactions as described in Agrawal *et al*., [36], and Bax, Chung, and Edge [37] and Dehghani-Tafti *et al*., [38]. However, how exactly the protons (hydrogens) move during catalysis is not yet clear (see below for details).

**Figure 5.**
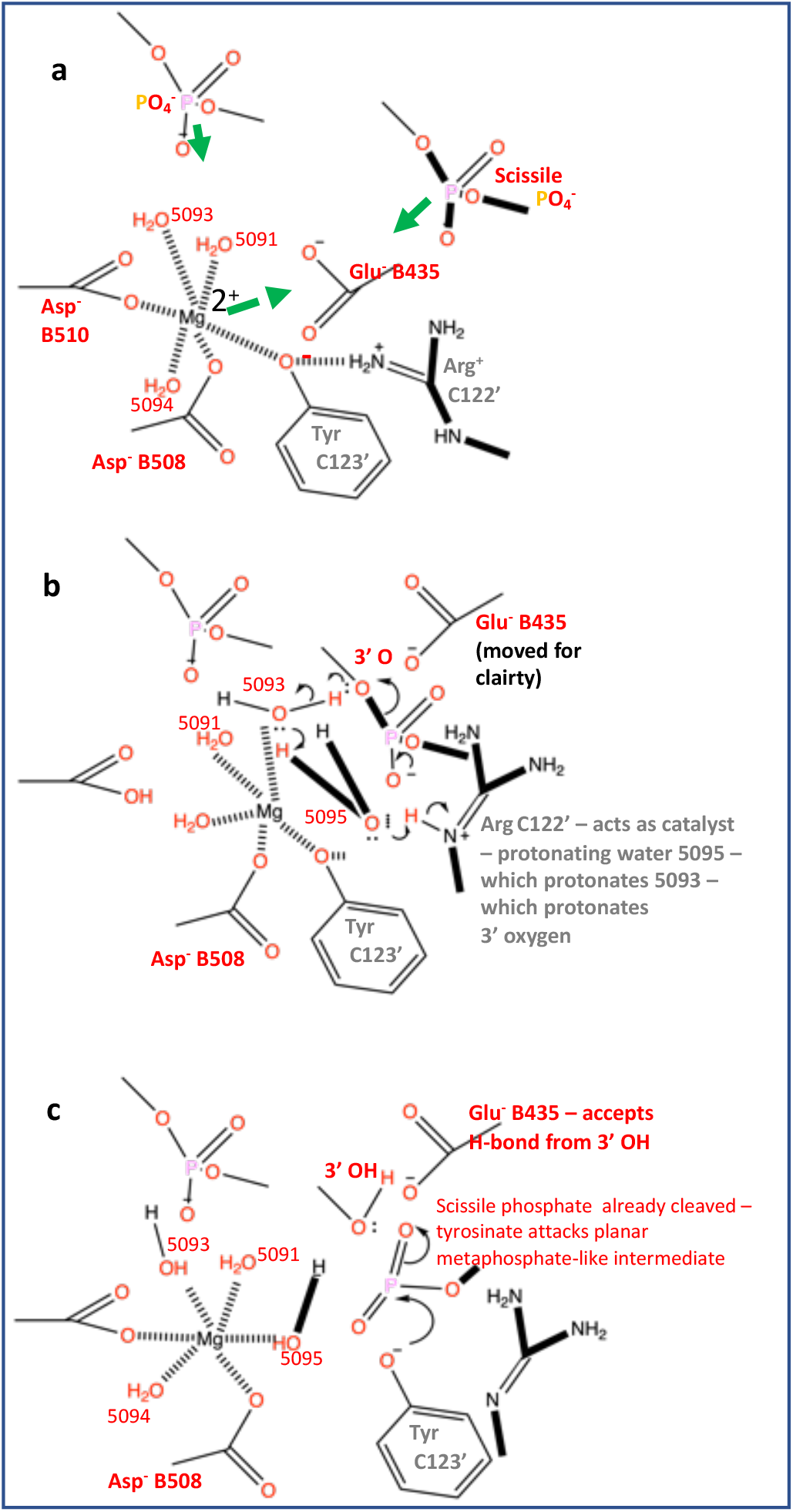
A mechanism for DNA-cleavage by *S.aureus* DNA gyrase. **(a)** The catalytic metal, Mg^2+^, comes into the active site pocket with the catalytic tyrosine (C123’). Because the side-chain oxygen is accepting a hydrogen bond from Arg 122’ and is also coordinating the e Mg^2+^ ion it is in the tyrosinate (negative ion) form already. The PO_4−_-ion prior to the scissile phosphate moves down to accept a hydrogen bond from water 5093. The Mg^2+^ ion is attracted towards Glu B435 and the scissile phosphate, and these are attracted towards the Mg^2+^ ion. **(b)** The Mg^2+^ ion has moved towards the A-site – this causes water 5093 to move and protonate the 3’ oxygen from the scissile phosphate, which then becomes the 3’ OH. However, almost simultaneously water 5095 comes into a new ‘hole’ and is immediately protonated by the N-epsilon hydrogen from Arg 122’. The scissile phosphate becomes a highly reactive planar metaphosphate-like group. **(c)** The catalytic tyrosinate ion, C123’, attacks the planar metaphosphate-like intermediate to generate the phosphotyrosine. Note Arg C122’ still needs to pick up a proton to regain its +1 charge. The catalytic metal returns to the Y(B) site with water 5095. In the presence of a T-DNA the cleaved G-DNA gate will open to allow T-DNA passage, before religation of the gate DNA.

One mechanism proposed in this paper (Figure 5) starts with protonation of the negatively charged DNA backbone, which generates a highly reactive metaphosphate like intermediate. If the negatively charged tyrosinate ion is not positioned to attack, then the metaphosphate will presumably immediately re-attack the 3’-OH, regenerating the uncleaved DNA. This mechanism seems chemically more sensible than the conventional mechanism which starts with the tyrosinate ion making a nucleophilic attack on a negatively charged tetrahedral phosphate from the DNA and ends with protonation of the 3’ oxygen (see for example the mechanism discussed in [8]). Although the proposed mechanism seems chemically more sensible - it is not proven and it depends on hydrogen atom transfer. Alteratively (supplementary Figure S9) water 5093 could be directly protonated by Arg 122’ to effect the DNA-cleavage. Such a mechanism (supplementary Figure S9) suggest that the single-stranded DNA-cleavage seen with NBTIs such as GSK299423 [12] and gepotidacin [39] is likely due to incomplete religation at one of the two active sites. Seeing such chemistry happen will require carefully designed experiments where hydrogen atoms could observed, such as by neutron [40] or electron diffraction experiments. Advancements over the last 10 years in electron diffraction show that this technique has potential to better locate hydrogen atoms, and could in the future be used to understand this mechanism further [41]. Experimental definition of hydrogen atom positions is challenging, and perhaps this explains why QM/MM studies [42] are still used despite their obvious limitations [2, 3]. A more definitive answer awaits the development of experimental techniques that allow hydrogens to be clearly visualized in multiple macromolecular structures.

In a 1993 paper entitled: ‘*A general two-metal-ion mechanism for catalytic RNA*’ Steitz and Steitz [43], proposed a rather pedestrian mechanism for catalytic RNA along the lines of Francis Crick’s old central dogma of molecular biology [44], i.e. the simple mantra “*DNA makes DNA makes RNA makes Protein*” [45]. The theory developed in their paper is based on four factors that were observed in early structures of the rather rigid catalytic proteins determined in the 1960s [46]. However, they seemed to have not noticed that RNA ‘invented proteins’ to be flexible molecular machines that could enhance catalytic rates by using ‘directional movement to protonate specific oxygen atoms on phosphates to catalyze phosphotransferase reactions’ [37]. While Tom Steitz’s 2009 Nobel Prize winning talk was entitled “*From the structure and function of the ribosome to new antibiotics*” [45], the company he co-founded in 2000, Rib-X (which sought to exploit expertise in (ribosomal) structural biology and molecular modeling to design new antibiotics) was taken over by Melinta Therapeutics and their first marketed drug was the DNA gyrase-targeting quinolone delafloxacin.

In 2010 Schmidt *et al*., [25] proposed a modified version of the Steitz and Steitz theory in ‘*A novel and unified two-metal mechanism for DNA cleavage by type II and IA topoisomerases*’ [25]. However, this proposal was based on a poorly refined low-resolution structure that does not have sensible metal-ion coordination geometry at either of the two different active sites. This structure has been re-refined (<u>RR-3l4k.pdb</u>) to have chemically reasonable geometry and a single metal ion at each active site, consistent with single moving metal mechanisms [13]. Similarly a paper by Bax *et al*., 2019 [13], contains a Figure 7 entitled ‘Dynamic model for the cleavage religation cycle catalyzed by a single metal ion’, which lacked sensible chemistry. Although it is difficult to disprove mechanisms [47], this does not mean they are true. We assert that proposed mechanisms should be consistent with the current knowledge of metal ion coordination geometry [26, 27].

We believe that the most likely current explanation of the observed synergy [25, 48-50] between different metal ions in type IIA topoisomerases is that when one active site contains a Y(B) site metal the other uses a 3’(A) site metal. Presumably differences in the ionic radius of calcium [26] account for calcium’s tendency to give more DNA cleavage in the absence of a compound [51, 52]. Similarities between type IA and type IIA topoisomerases, which both have Toprim domains, suggest that they likely have similar mechanisms [8]. Although type IA topoisomerases only work on negatively supercoiled DNA and can cleave a single-stranded DNA segment in the absence of a divalent metal ion [8] they require a divalent metal ion for DNA-religation [8, 53]. Dual targeting of DNA gyrase and topoisomerase IV [54] is important to prevent the likelihood of developing resistance to drugs. If a compound could be found that inhibits both *M. tb* topo I and DNA gyrase (Supplementary Figure S1) it would be interesting to measure the likelihood of resistance to such a compound.

## 4. Materials and Methods

The aim of this paper is to present a new moving metal mechanism which is consistent with existing crystal structures. The mechanism depends on the accuracy of the metal-ion coordination geometry and re-refined coordinates are available [11] for the two crystal structures containing catalytic metal ions (<u>2xcs-v2-BA-x.pdb</u> and <u>5cdm-v2-BA-x.pdb</u>). However, we note that crystal structures are static pictures of thousands of molecules that make up a crystal.

What we are concerned with in this paper is how DNA-gyrase cleaves DNA. To do this we used the following procedure: (i) Coordinates from crystal structures were manually edited in coot [55] - so that each coordinate set contained a complete set of single atoms. Also the DNA sequence was manually edited so that it was the same in the three starting structures (see Table 2 for details). This gave: 2xcs-v2-one-DNA.pdb, 5cdm-v2-one-DNA.pdb and 6fqv-c1-one-DNA.pdb. Note 6fqv-c1-one-DNA.pdb contains no catalytic metal ion and Glu B435 was modelled in in a conformation similar to that observed in 5cdm. The two sets of coordinates with a metal ion, 2xcs-v2-one-DNA.pdb, 5cdm-v2-one-DNA.pdb, were read into Maestro [32] and; (ii) hydrogens were added onto all atoms in ‘prepare’ [56]. The structure was written out from Maestro (export structure) and edited so the header contained a CRYST1 record. Then, because the hydrogen atoms placed on waters were all pointing in the same direction, the relevant waters were manually rotated (in coot [55]) around the oxygen atom to chemically sensible positions [57]. The coordinates with manually rotated waters were then read back into Maestro [32], the relevant waters, including hydrogens were selected and then energy minimized in Maestro (minimize selected atoms). The manganese atom had a formal charge of +2 in Maestro. We used this procedure to keep heavy atom coordinates (non-hydrogen atoms) used for making movies close to those experimentally defined from crystal structures [11]. This gave: 2xcs-v2-two-DNA.pdb, 5cdm-v2-two-DNA.pdb as explained in Table 2; and with and 6fqv-c1-one.pdb, these were used to help guide the making of the molecular DNA-cleavage movie

To aid visualisation of our proposed mechanism of DNA cleavage in type IIA topoisomerases, we created a molecular movie that morphs between four states (Table 3): (i) uncleaved DNA with no metal ion present; (ii) entry of the metal ion coordinated by the catalytic tyrosine; (iii) transient state at the point of DNA cleavage; and (iv) cleaved DNA with the metal ion still bound. The movie was produced using the ProSMART library for structure analysis [58], similarly as in [59]. The procedure involves applying hierarchical aggregate transformations to residues, backbone and side chain atoms, resulting in a parsimonious morphing between states. This required preparation of coordinate models corresponding to four states.

State 1 - uncleaved DNA (6fqv-c2): Due to this model being the second complex in 6fqv, the chain nomenclature is different due to the requirement for uniqueness. Specifically, the protein chains named R/S/T/U (authors’ original annotation) correspond to the conventionally named chains A/B/C/D, and similarly the nucleic acid chains V/W correspond to E/F. To achieve an equivalent coordinate frame for producing the movie, the complex was superposed onto the other complex, aligning chain S in 6fqv-c2 to chain B in 6fqv-c1 using ProSMART [58]. In 6fqv-c2, Arg122 is modelled in two conformations; conformer A was removed and B retained, so as to retain the hydrogen bond between the Tyr C123 terminal oxygen and Arg C122(NH2); an interaction that is relevant to the proposed mechanism.

State 2 - entry of the metal ion (6fqv-c1): Despite the metal ion not being present in 6fqv, we transplanted the Mn^2+^ and coordinating waters from 5cdm, keeping the metal ion coordinated to Asp B508(OD2) and Asp B510(OD2). We manually adjusted the positions of the coordinating waters (B5091, B5093 and B5094) to ensure reasonable coordination geometry, with hydrogen positions optimised using Maestro [32]. Model 5cdm also has water B5095 coordinating the metal. However, we propose that the metal is brought into the binding site by Tyr C123, and that the C123 side chain oxygen atom takes the place of the water B5095 observed in 5cdm. Consequently, we manually adjusted the position of Tyr C123 (by rotating the side chain) so that the C123 side chain oxygen atom coordinates the metal ion in place of B5095. This resulted in the C123 side chain oxygen atom moving a distance of some 1.3 Å. Similarly, we manually adjusted the adjacent Arg C122 so as to maintain the hydrogen bond between C122(NH2) and side chain oxygen atom of C123. These ‘manual’ adjustments were applied in Coot [55], ensuring reasonable geometry of the adjusted residues.

State 3 - transient state at the point of DNA cleavage (between 6fqv-c1 and 2xcs): The model 2xcs exhibits uncleaved DNA, with the catalytic tyrosine C123 mutated to phenylalanine, and the metal ion coordinated by the OP1 atom of the uncleaved phosphate DG E9. Our proposed mechanism involves the metal ion inducing cleavage prior to the structure entering the state observed in 2xcs - i.e. it is proposed that the metal ion does not quite reach the scissile phosphate. However, we do not have access to an intermediate state at the exact point at which cleavage occurs; it is clear why it would be difficult to observe such a transient state using conventional crystallographic methods. Consequently, it was necessary to generate an intermediate between the previous state (derived from 6fqv-c1) and 2xcs. This State 3 was generated by morphing between State 2 and 2xcs (using ProSMART, as for the main movie), and extracting an intermediate set of coordinates at the point at which (we hypothesise) the DNA-cleavage-inducing chemical reaction occurs.

We also hypothesized that a water moves into the binding site, to a position (reasonably close to water E52 in 2xcs) that is between Arg C122, Tyr C123, water B5093 and the phosphate of DG9. We manually placed such a water in a position that we believed most chemically sensible. In the movie, this water is seen to move in through an open pocket just prior to DNA cleavage. This water (E52=5095 in Figure 5 and Nicholls-et-al-movie1.mp4) is protonated by Arg 122’ in Figure 5 and the movie. However, as can be seen if figure 3, if Arg 122’ moves more slowly another possibility is that Arg 122’ directly protonates water 5093 (see supplementary Figure S9).

State 4 - cleaved DNA (5cdm): The final state is that with cleaved DNA observed in 5cdm.

## 5. Conclusions

In 1979 Brown and Cozarelli [60] proposed a sign inversion mechanism to explain the observed activity of *E.coli* DNA gyrase. They proposed that: ‘*a positive supercoil is directly inverted into a negative one via a transient doublestrand break*’ and that the ‘*reverse process is the path by which gyrase, in the absense of ATP, relaxes negatively supercoiled DNA*.’ Since the initially proposed model [30] was based on relaxation of negatively supercoiled DNA (and decatenation), it seems logical to suggest that relaxation of positively supercoiled DNA should proceed by the reverse direction of strand transfer (see Figure 6). If this is the case then the small Greek Key domains would be ideally placed to regulate access of metal ions to the active sites for fast relaxation of positively supercoiled DNA.

**Figure 6.**
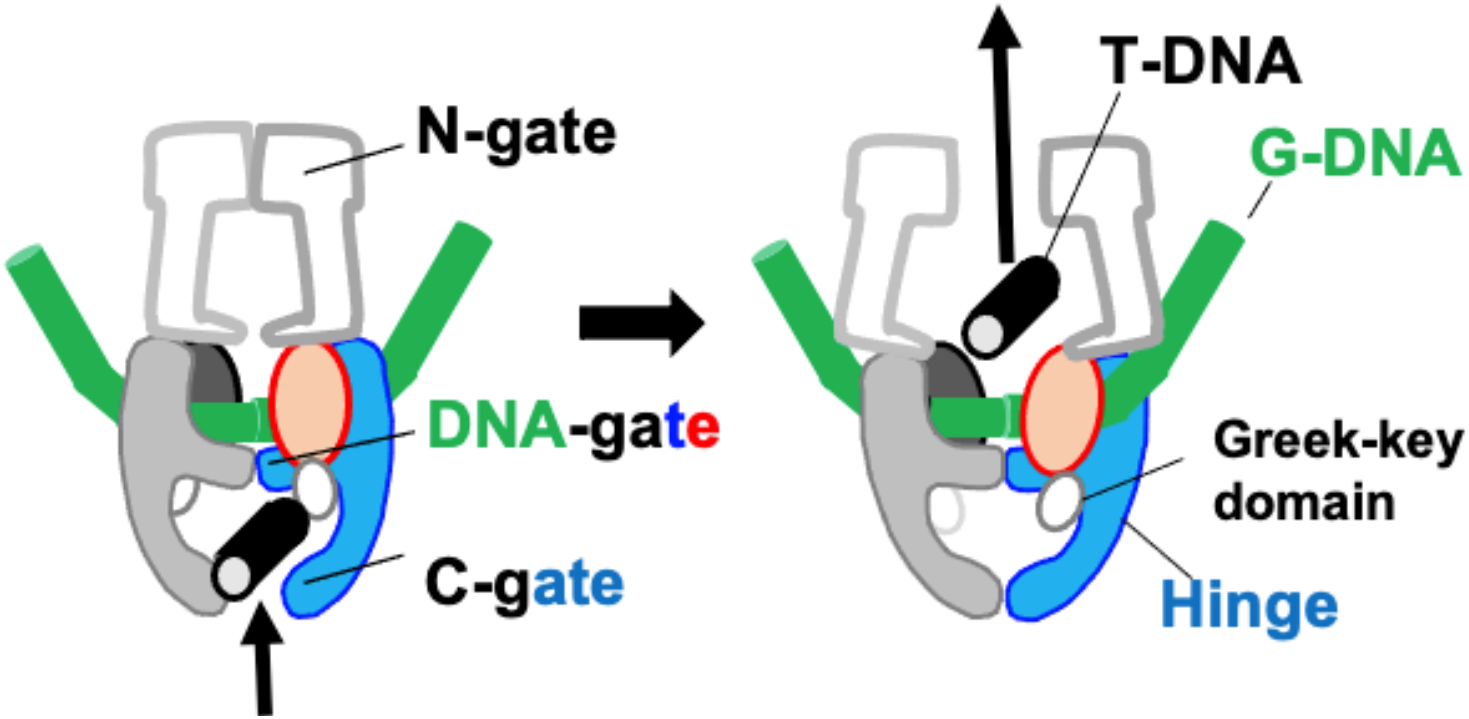
Could the T-DNA move ‘up’ when relaxing positively supercoiled DNA? The C-terminal DNA wrapping domains have been omitted for simplicity

Human topoisomerase IIα and DNA gyrase are able to relax positively supercoiled DNA at enhanced rates ahead of replication forks in humans [21, 22] and bacteria [23, 24]. A simple model would be that the N(ATP-dependent)-gate tends to be closed during relaxation of positive supercoils; i.e. that the enzyme acts rather like a rocker switch with the ATPase and G-gates closed for relaxation of positive supercoils, but the C-gate open. In such a simplistic model the C-terminal domains of human topoisomerase IIα or DNA gyrase would bind positively supercoiled plectonemes in front of replication forks [61], and the small Greek Key domains (Figures 6 and 7) would recognise the incoming T-DNA segments facilitating rapid and safe relaxation of positive supercoils [62] in front of replication forks.

**Figure 7.**
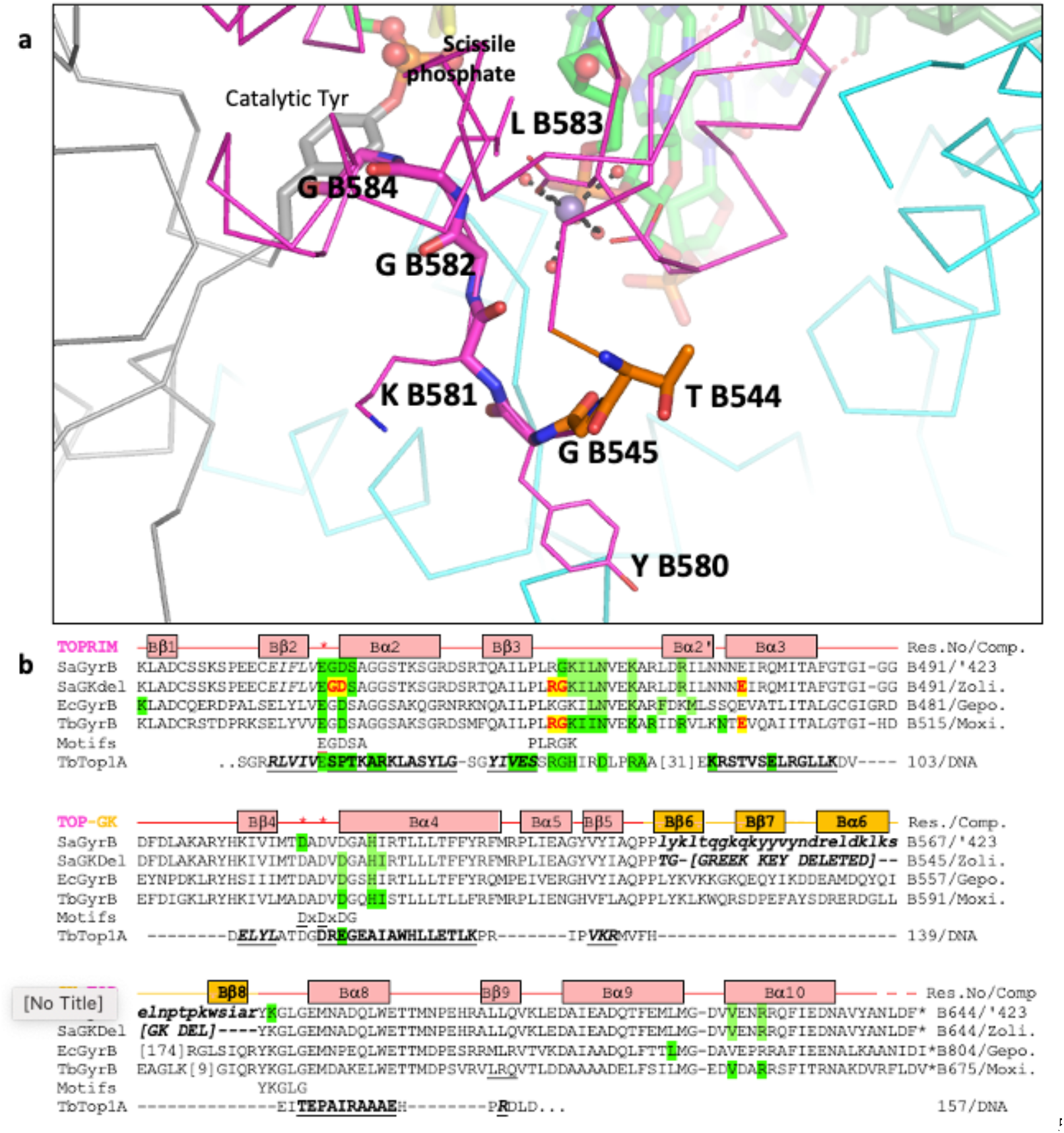
The Greek Key deletion and YKGLG motif are close to the catalytic site. **(a)** A view of the position the two residues, T B544 and G B545 (orange carbons), that have replaced the ‘Greek Key’ domain in 5cdm-BA-x.pdb. Note the Greek Key motif, named after a pattern that was common on Greek pottery, normally contains four β-strands connected by hairpins (https://www.cryst.bbk.ac.uk/PPS2/course/section10/all_beta.html). In type IIA topoisomerases the third β-strand of the Greek Key is replaced by an alpha helix and insertions in Gram negative DNA gyrases are found in the connection to the final β-strand. Because the final β-strand is adjacent to the first β-strand it has been possible to delete the Greek Key domain in *S. aureus* DNA gyrase. **(b)** an alignment of the amino acid sequences from four Gyrase structures: *S.aureus* GyrB + GSK299423 (SaGyrB 3.5Å - PDB code: 2XCR), *S.aureus* GyrB + zoliflodacin (SaGKdel 2.8Å - 8BP2), *E. coli* GyrB + gepotidacin (EcGyrB 4.0Å - 6RKS), and *M. tuberculosis* GyrB + moxifloxacin (TbGyrB 2.4Å - 5BS8). The *S. aureus* structures are crystal structures of the catalytic core (see figure 1) the *E.coli* structure is a cryoEM structure containing full length GyrA and GyrB. The motifs line highlights four conserved sequence motifs, three of which: (i) <u>E</u>GDSA, and (ii) <u>D</u>x<u>D</u>xDG and (iii) the YKGLG motif (after the greek key domain) are close to the catalytic metal. Amino acids are highlighted on the sequences if they contact (< 3.8 Å) either the **compound**, or the DNA (green); (note zoliflodacin **GD** and **RG** residues also contact the DNA; only the **G** from the **RG** motif also contacts the DNA in the moxifloxacin structure). The *M. tuberculosis* Top1A (TbTop1A) sequence is from the 2.78Å structure with T and G-segment DNA (PDB code: 8CZQ); only the Toprim domain sequence is shown with secondary structural elements underlined in bold; the GK-TOP line shows the secondary structural elements from the 3.5Å SaGyrB structure with GSK299423. Numbers indicate extra residues not shown (e.g.[31]). (see supplementary Figure 5 in Chan *et al*., 2015 for contacts on GyrA).

While the the ATPase domains are not 100% required for relaxation of negative supercoils by human topoisomerase IIβ [63] similar experiments have not yet been done, to the best of our knowledge, for relaxation of positive supercoils. The generation of positively supercoiled plasmids with reverse gyrase [64] may be technically more challenging than the generation of negatively supercoiled plasmids with DNA gyrase, however, if Brown and Cozzarelli [60] are correct then the relaxation of positively supercoiled DNA may proceed in the opposite direction from the relaxation of negatively supercoiled DNA; such experiments may be worthwhile.

## Supplementary Materials

The following supporting informationis available, Figure S1: A drug design relevant classification of topoisomerases. Figure S2: Orthogonal views of three moxifloxacin DNA-cleavage complexes. Figure S3: Orthogonal views of a moxifloxacin DNA-cleavage complex. Supplementary Figure S4. Sequence alignment of GyrB from M. tuberculosis (GYRB_MYCTU), E. coli and S. aureus. Figure S5: Catalytic Mn2^+^ ions in crystal structures of *S. aureus* DNA gyrase. Supplementary Figure S6. Sequence alignment of GyrA from M. tuberculosis (GYRB_MYCTU), E. coli, S. aureus and B. subtillis. (in two parts) Figure S7: The presence of an Mn2+ ion at the Y(B) site seems to order Glu B435 and attract the phosphate before the scissile phosphate towards water 5093. Figure S8: A comparison of 6fqv-c1-one-DNA.pdb with 2xcs-v2-two-DNA.pdb and 5cdm-v2-two-DNA.pdb. Supplementary Figure S9. Could water 5093 have momentarily cleaved the DNA?. and **Video S1: Nicholls-et-al-movie1.mp4**

## Author Contributions

“Conceptualization, B.D.B. and R.A.N.; methodology, B.D.B. and R.A.N; software, R.A.N., F.L and G.A.N.; validation, B.D.B., R.A.N., F.L and G.A.N.; formal analysis, B.D.B, H.M., D.S. and R.A.N.; investigation, R.A.N., H.M., D.S. and B.D.B.; resources, R.A.N., A.J.W and S.W.; data curation, B.D.B.; writing—original draft preparation, B.D.B.; writing—review and editing, All authors.; visualization, B.D.B., D.S. and R.A.N.; supervision, B.D.B, A.J.W. and S.W..; project administration, R.A.N., B.D.B, A.J.W. and S.W.; funding acquisition, R.A.N., A.J.W and S.W.. All authors have read and agreed to the published version of the manuscript.” Please turn to the <u>CRediT taxonomy</u> for the term explanation. Authorship must be limited to those who have contributed substantially to the work reported.

## References

1. Raithby, P.R., Single Crystal and Powder Methods for Structure Determination of Metastable Species, in The Future of Dynamic Structural Science. 2013, Springer. p. 1–12.

2. Musielak, Z.E. and B. Quarles, The three-body problem. Reports on Progress in Physics, 2014. 77(6): p. 065901.

3. Belbruno, E., Relation between solutions of the Schrödinger equation with transitioning resonance solutions of the gravitational three-body problem. Journal of Physics Communications, 2020. 4(1): p. 015012.

4. Schoeffler, A.J. and J.M. Berger, DNA topoisomerases: harnessing and constraining energy to govern chromosome topology. Q. Rev. Biophys, 2008. 41(1): p. 41–101.

5. Bates, A.D. and A. Maxwell, DNA topology. 2005: Oxford University Press, USA.

6. Pommier, Y., et al., Roles of eukaryotic topoisomerases in transcription, replication and genomic stability. Nat Rev Mol Cell Biol, 2016. 17(11): p. 703–721.

7. Ahmed, W., et al., Conditional silencing of topoisomerase I gene of Mycobacterium tuberculosis validates its essentiality for cell survival. FEMS microbiology letters, 2014. 353(2): p. 116–123.

8. Dasgupta, T., S. Ferdous, and Y.C. Tse-Dinh, Mechanism of Type IA Topoisomerases. Molecules, 2020. 25(20).

9. Vayssières, M., et al., Structural basis of DNA crossover capture by Escherichia coli DNA gyrase. Science, 2024. 384(6692): p. 227–232.

10. Michalczyk, E., et al., Structure of Escherichia coli DNA gyrase with chirally wrapped DNA supports ratchet-and-pawl mechanism for an ATP-powered supercoiling motor. bioRxiv, 2024: p. 2024.04. 12.589215.

11. Morgan, H., et al., How do gepotidacin and zolifodacin stabilize DNA-cleavage complexes with bacterial type IIA topoisomerases? 1. Experimental definition of metal binding sites. Bioarchives, 2024.

12. Bax, B.D., et al., Type IIA topoisomerase inhibition by a new class of antibacterial agents. Nature, 2010. 466(7309): p. 935–940.

13. Bax, B.D., et al., DNA Topoisomerase Inhibitors: Trapping a DNA-Cleaving Machine in Motion. J Mol Biol, 2019. 431(18): p. 3427–3449.

14. Wohlkonig, A., et al., Structural basis of quinolone inhibition of type IIA topoisomerases and target-mediated resistance. Nat. Struct. Mol. Biol, 2010. 17(9): p. 1152–1153.

15. Aldred, K.J., et al., Topoisomerase IV-quinolone interactions are mediated through a water-metal ion bridge: mechanistic basis of quinolone resistance. Nucleic Acids Res, 2013. 41(8): p. 4628–4639.

16. Laponogov, I., et al., Structural insight into the quinolone-DNA cleavage complex of type IIA topoisomerases. Nat. Struct. Mol. Biol, 2009. 16(6): p. 667–669.

17. Blower, T.R., et al., Crystal structure and stability of gyrase-fluoroquinolone cleaved complexes from Mycobacterium tuberculosis. Proc. Natl. Acad. Sci. U. S. A, 2016. 113(7): p. 1706–1713.

18. Chan, P.F., et al., Structural basis of DNA gyrase inhibition by antibacterial QPT-1, anticancer drug etoposide and moxifloxacin. Nat. Commun, 2015. 6: p. 10048.

19. Srikannathasan, V., et al., Crystallization and initial crystallographic analysis of covalent DNA-cleavage complexes of Staphyloccocus aureus DNA gyrase with QPT-1, moxifloxacin and etoposide. Acta Crystallographica Section F: Structural Biology Communications, 2015. 71(10): p. 1242–1246.

20. Pommier, Y., et al., Human topoisomerases and their roles in genome stability and organization. Nature Reviews Molecular Cell Biology, 2022. 23(6): p. 407–427.

21. McClendon, A.K., A.C. Rodriguez, and N. Osheroff, Human topoisomerase IIalpha rapidly relaxes positively supercoiled DNA: implications for enzyme action ahead of replication forks. J. Biol. Chem, 2005. 280(47): p. 39337–39345.

22. McClendon, A.K., et al., Bimodal recognition of DNA geometry by human topoisomerase IIα: preferential relaxation of positively supercoiled DNA requires elements in the C-terminal domain. Biochemistry, 2008. 47(50): p. 13169–13178.

23. Ashley, R.E., et al., Activities of gyrase and topoisomerase IV on positively supercoiled DNA. Nucleic acids research, 2017. 45(16): p. 9611–9624.

24. Ashley, R.E., et al., Recognition of DNA supercoil geometry by Mycobacterium tuberculosis gyrase. Biochemistry, 2017. 56(40): p. 5440–5448.

25. Schmidt, B.H., et al., A novel and unified two-metal mechanism for DNA cleavage by type II and IA topoisomerases. Nature, 2010. 465(7298): p. 641–644.

26. Katz, A.K., et al., Calcium ion coordination: a comparison with that of beryllium, magnesium, and zinc. Journal of the American Chemical Society, 1996. 118(24): p. 5752–5763.

27. Bock, C.W., et al., Manganese as a Replacement for Magnesium and Zinc: Functional Comparison of the Divalent Ions. J. Am. Chem. Soc, 1999. 121: p. 7360–7372.

28. Germe, T., et al., A new class of antibacterials, the imidazopyrazinones, reveal structural transitions involved in DNA gyrase poisoning and mechanisms of resistance. Nucleic Acids Res, 2018. 46(8): p. 4114–4128.

29. Aravind, L., D.D. Leipe, and E.V. Koonin, Toprim--a conserved catalytic domain in type IA and II topoisomerases, DnaG-type primases, OLD family nucleases and RecR proteins. Nucleic Acids Res, 1998. 26(18): p. 4205–4213.

30. Berger, J.M., et al., Structure and mechanism of DNA topoisomerase II. Nature, 1996. 379(6562): p. 225–232.

31. Wu, C.C., et al., Structural basis of type II topoisomerase inhibition by the anticancer drug etoposide. Science, 2011. 333(6041): p. 459–462.

32. Maestro, S., Maestro. Schrödinger, LLC, New York, NY, 2020. 2020: p. 682.

33. The PyMOL Molecular Graphics System, Version 1.5.0.4 Schrödinger, LLC. 2013.

34. Changela, A., R.J. DiGate, and A. Mondragón, Crystal structure of a complex of a type IA DNA topoisomerase with a single-stranded DNA molecule. Nature, 2001. 411(6841): p. 1077–1081.

35. Frey, P.A. and A.D. Hegeman, Enzymatic Reaction Mechanisms. 2007, New York: Oxford University Press.

36. Agrawal, A., et al., Mycobacterium tuberculosis DNA gyrase ATPase domain structures suggest a dissociative mechanism that explains how ATP hydrolysis is coupled to domain motion. Biochem J, 2013. 456(2): p. 263–73.

37. Bax, B., C.W. Chung, and C. Edge, Getting the chemistry right: protonation, tautomers and the importance of H atoms in biological chemistry. Acta Crystallogr D Struct Biol, 2017. 73(Pt 2): p. 131–140.

38. Dehghani-Tafti, S., et al., Structural and functional analysis of the nucleotide and DNA binding activities of the human PIF1 helicase. Nucleic Acids Res, 2019. 47(6): p. 3208–3222.

39. Gibson, E.G., et al., Mechanistic and Structural Basis for the Actions of the Antibacterial Gepotidacin against Staphylococcus aureus Gyrase. ACS Infect Dis, 2019. 5(4): p. 570–581.

40. Liebschner, D., et al., Implementation of the riding hydrogen model in CCTBX to support the next generation of X-ray and neutron joint refinement in Phenix. Methods Enzymol, 2020. 634: p. 177–199.

41. Clabbers, M.T., et al., Hydrogens and hydrogen-bond networks in macromolecular MicroED data. Journal of Structural Biology: X, 2022. 6: p. 100078.

42. Berta, D., et al., Cations in motion: QM/MM studies of the dynamic and electrostatic roles of H+ and Mg2+ ions in enzyme reactions. Current Opinion in Structural Biology, 2020. 61: p. 198–206.

43. Steitz, T.A. and J.A. Steitz, A general two-metal-ion mechanism for catalytic RNA. Proc Natl Acad Sci U S A, 1993. 90(14): p. 6498–502.

44. Crick, F., Central dogma of molecular biology. Nature, 1970. 227(5258): p. 561–563.

45. Steitz, T.A., From the structure and function of the ribosome to new antibiotics (Nobel Lecture). Angewandte Chemie International Edition, 2010. 49(26): p. 4381–4398.

46. Blow, D. and T. Steitz, X-ray diffraction studies of enzymes. Annual review of Biochemistry, 1970. 39(1): p. 63–100.

47. Wlodawer, A., et al., Detect, correct, retract: How to manage incorrect structural models. The FEBS journal, 2018. 285(3): p. 444–466.

48. Deweese, J.E., A.B. Burgin, and N. Osheroff, Human topoisomerase IIalpha uses a two-metal-ion mechanism for DNA cleavage. Nucleic Acids Res, 2008. 36(15): p. 4883–4893.

49. Deweese, J.E., et al., Metal ion interactions in the DNA cleavage/ligation active site of human topoisomerase IIalpha. Biochemistry, 2009. 48(38): p. 8940–8947.

50. Deweese, J.E. and N. Osheroff, The use of divalent metal ions by type II topoisomerases. Metallomics, 2010. 2(7): p. 450–459.

51. Osheroff, N. and E.L. Zechiedrich, Calcium-promoted DNA cleavage by eukaryotic topoisomerase II: trapping the covalent enzyme-DNA complex in an active form. Biochemistry, 1987. 26(14): p. 4303–4309.

52. Aldred, K.J., et al., Fluoroquinolone interactions with Mycobacterium tuberculosis gyrase: Enhancing drug activity against wild-type and resistant gyrase. Proc Natl Acad Sci U S A, 2016. 113(7): p. E839–46.

53. Narula, G., et al., The strictly conserved Arg-321 residue in the active site of Escherichia coli topoisomerase I plays a critical role in DNA rejoining. Journal of Biological Chemistry, 2011. 286(21): p. 18673–18680.

54. Oviatt, A.A., et al., Interactions between Gepotidacin and Escherichia coli Gyrase and Topoisomerase IV: Genetic and Biochemical Evidence for Well-Balanced Dual-Targeting. ACS Infectious Diseases, 2024.

55. Emsley, P., et al., Features and development of Coot. Acta Crystallogr. D. Biol. Crystallogr, 2010. 66(Pt 4): p. 486–501.

56. Shelley, J.C., et al., Epik: a software program for pK a prediction and protonation state generation for drug-like molecules. Journal of computer-aided molecular design, 2007. 21: p. 681–691.

57. Jeffrey, G.A. and W. Saenger, Hydrogen bonding in biological structures. 2012: Sprimger Science & Business Media.

58. Nicholls, R.A., et al., Conformation-independent structural comparison of macromolecules with ProSMART. Acta Crystallographica Section D: Biological Crystallography, 2014. 70(9): p. 2487–2499.

59. Koulouris, C.R., et al., Tyrosine 121 moves revealing a ligandable pocket that couples catalysis to ATP-binding in serine racemase. Communications Biology, 2022. 5(1): p. 346.

60. Brown, P.O. and N.R. Cozzarelli, A sign inversion mechanism for enzymatic supercoiling of DNA. Science, 1979. 206(4422): p. 1081–1083.

61. Skoruppa, E. and E. Carlon, Equilibrium fluctuations of DNA plectonemes. Physical Review E, 2022. 106(2): p. 024412.

62. Zechiedrich, E.L. and N. Osheroff, Eukaryotic topoisomerases recognize nucleic acid topology by preferentially interacting with DNA crossovers. The EMBO journal, 1990. 9(13): p. 4555–4562.

63. Bandak, A.F., et al., Using energy to go downhill—a genoprotective role for ATPase activity in DNA topoisomerase II. Nucleic acids research, 2024. 52(3): p. 1313–1324.

64. Kikuchi, A. and K. Asai, Reverse gyrase—a topoisomerase which introduces positive superhelical turns into DNA. Nature, 1984. 309(5970): p. 677–681.

