## Supplementary figures for "How do gepotidacin and zolifodacin stabilize DNA-cleavage complexes with bacterial type IIA topoisomerases? 2. A Single Moving Metal Mechanism"

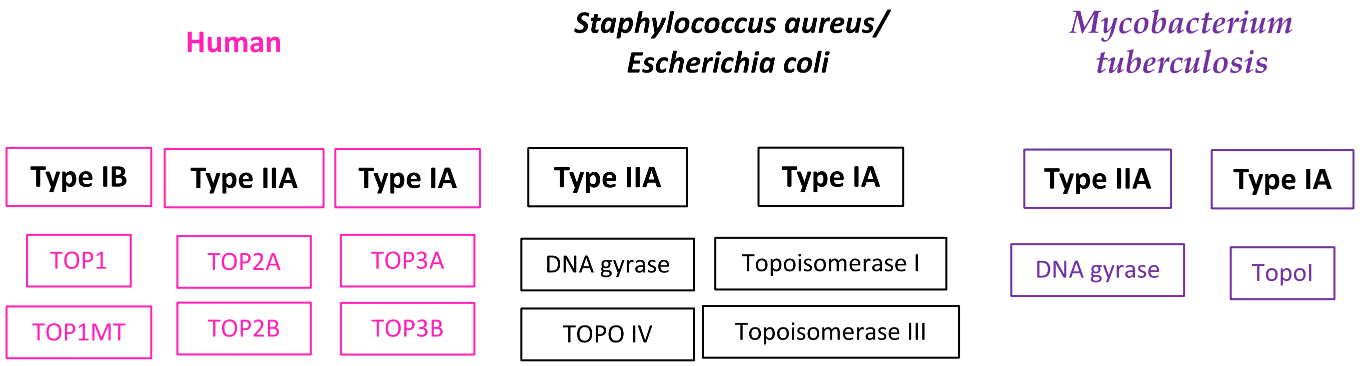


**Supplementary Figure S1: A drug design relevant classification of topoisomerases.** ‘*Human topoisomerases comprise a family of six enzymes: two type IB (TOP1 and mitochondrial TOP1 - TOP1MT), two type IIA (TOP2A and TOP2B) and two type IA (TOP3A and TOP3B) topoisomerases*’ (Pommier, Y., et al., Human topoisomerases and their roles in genome stability and organization. Nature Reviews Molecular Cell Biology, 2022. 23(6): p. 407-427). The two type IB enzymes cleave the DNA creating a 3′-phosphodiester bond with the catalytic tyrosine; the type IIA and type IA topoisomerases cleave the DNA using a Toprim domain and create a 5′-phosphodiester bond with the catalytic tyrosine. Interestingely 'the Mtb genome encodes only one copy of type I and one copy of type II topoisomerase’ (Ahmed, W., *et al.*, Conditional silencing of topoisomerase I gene of Mycobacterium tuberculosis validates its essentiality for cell survival. FEMS microbiology letters, 2014. 353(2): p. 116-123). Type IIB topoisomerases are not current drug targets and are therefore not shown. Type IIB topoisomerases seem to be involved in meiosis, playing a role in genetic recombination (in which DNA is ‘cut and resectioned’).

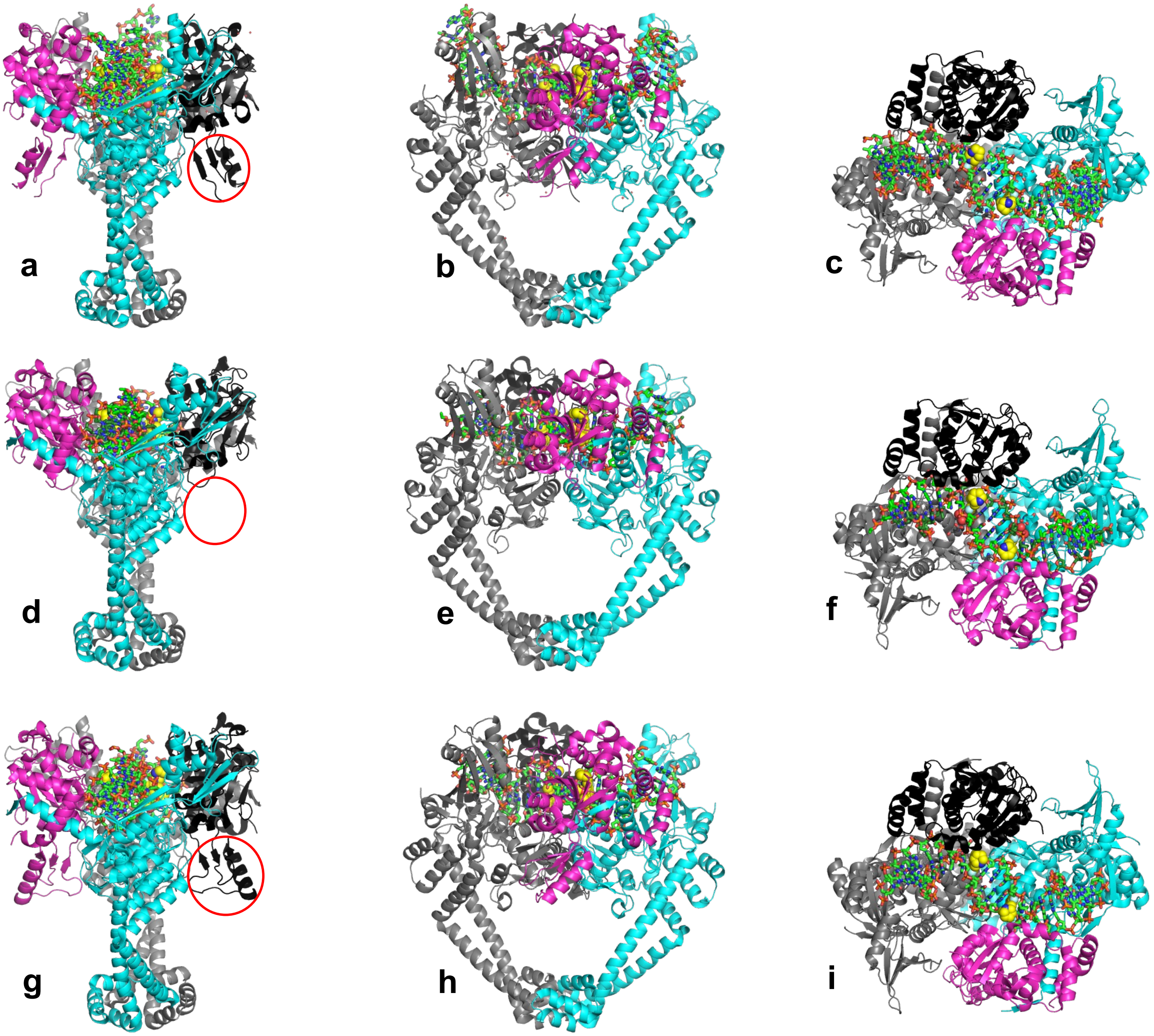


**Supplementary Figure S2. Orthogonal views of three moxifloxacin DNA-cleavage complexes.**
**(a,b,c)** Orthogonal (90º) views of the 3.25Å *A. baumanii* topoisomerase IV crystal structure (pdb code: 2xkk). ParC subunits coloured cyan and grey. ParE subunits magenta and black. Note the presence of the small Greek Key domain seen in view a (red circle on black ParE subunit). The two moxifloxacin molecules are shown in sphere representation with carbons in yellow.
**(d,e,f)** Orthogonal views of the 2.95Å *S. aureus* DNA gyrase crystal structure (pdb code: 5cdq). GyrA subunits coloured cyan and grey. GyrB subunits magenta and black. Note the absence of the small Greek Key domain seen in view d. The two moxifloxacin molecules are shown in sphere represenation with carbons in yellow.
**(g,h,i)** Orthogonal views of the 2.4Å *M. tuberculosis* DNA gyrase crystal structure (pdb code: 5bs8). GyrA subunits coloured cyan and grey. GyrB subunits magenta and black. Note the presence of the small Greek Key domain seen in view g (red circle on black GyrB subunit). The two moxifloxacin molecules are shown in sphere represenation with carbons in yellow. For clarity hydrogens are not shown.

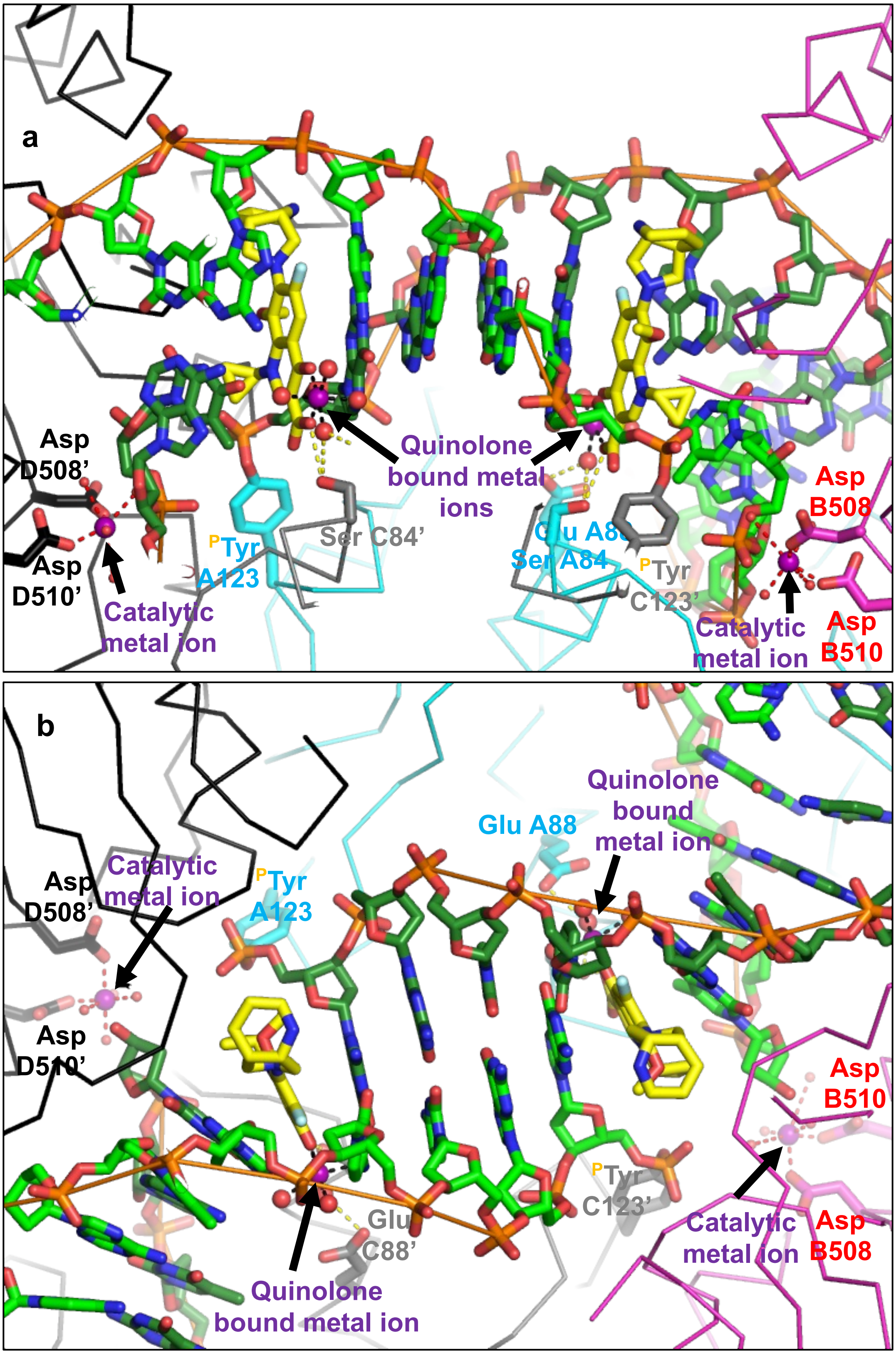


**Supplementary Figure S3. Orthogonal views of a moxifloxacin DNA-cleavage complex.**
**(a)** View of the 2.95Å *S. aureus* DNA gyrase crystal structure (pdb code: 5cdq). GyrA subunits coloured cyan and grey, shown as Cα ‘ribbon’ , with labeled side-chains shown in ‘stick’. GyrB subunits magenta and black. The two moxifloxacin molecules are shown in stick representation with carbons in yellow; cleaved DNA carbons in green.
**(b)** Orthogonal (90º) view of the same structure.

```
GYRB_MYCTU      --MAAQKKKAQDEYGAASITILEGLEAVRKRPGMYIGSTG-ERGLHHLIWEVVDNAVDEA 57
GYRB_ECOLI      ---------MSNSYDSSSIKVLKGLDAVRKRPGMYIGDTDDGTGLHHMVFEVVDNAIDEA 51
GYRB_STAAN      MVTALSDVNNTDNYGAGQIQVLEGLEAVRKRPGMYIGSTS-ERGLHHLVWEIVDNSIDEA 59
                           :.*.:..* :*:**:***********.*.   ****:::*:***::***

GYRB_MYCTU      MAGYATTVNVVLLEDGGVEVADDGRGIPVATHA-SGIPTVDVVMTQLHAGGKFDSDAYAI 116
GYRB_ECOLI      LAGHCKEIIVTIHADNSVSVQDDGRGIPTGIHPEEGVSAAEVIMTVLHAGGKFDDNSYKV 111
GYRB_STAAN      LAGYANKIEVVIEKDNWIKVTDNGRGIPVDIQEKMGRPAVEVILTVLHAGGKFGGGGYKV 119
                :**:.. : *.:   *. :.* *:*****.   :    *   :.:*::* *******....* :

GYRB_MYCTU      SGGLHGVGVSVVNALSTRLEVEIKRDGYEWSQVYEKSEPL-GLKQGAPTKKTGSTVRFWA 175
GYRB_ECOLI      SGGLHGVGVSVVNALSQKLELVIQREGKIHRQIYEHGVPQAPLAVTGETEKTGTMVRFWP 171
GYRB_STAAN      SGGLHGVGSSVVNALSQDLEVYVHRNETIYHQAYKKGVPQFDLKEVGTTDKTGTVIRFKA 179
                ******** *******   **: ::*:      * *::. *    *    . *.***: :**

GYRB_MYCTU      DPAVF-ETTEYDFETVARRLQEMAFLNKGLTINLTDERVTQDEVVDEVVSDVAEAPKSAS 234
GYRB_ECOLI      SLETFTNVTEFEYEILAKRLRELSFLNSGVSIRLRDKRDG-------------------- 211
GYRB_STAAN      DGEIFTETTVYNYETLQQRIRELAFLNKGIQITLRDERDE-------------------- 219
                .   * :.* :::* : :*::*::***.*: * * *:*

GYRB_MYCTU      ERAAESTAPHKVKSRTFHYPGGLVDFVKHINRTKNAIHSSIVDFSGKGTGHEVEIAMQWN 294
GYRB_ECOLI      ------------KEDHFHYEGGIKAFVEYLNKNKTPIHPNIFYFSTEKDGIGVEVALQWN 259
GYRB_STAAN      ---------ENVREDSYHYEGGIKSYVELLNENKEPIHDEPIYIHQSKDDIEVEIAIQYN 270
                            :.   :** **:   :*: :*..*   ** . . :   .   .  **:*:*:*

GYRB_MYCTU      AGYSESVHTFANTINTHEGGTHEEGFRSALTSVVNKYAKDRKLLKDKDPNLTGDDIREGL 354
GYRB_ECOLI      DGFQENIYCFTNNIPQRDGGTHLAGFRAAMTRTLNAYMDKEGYSKKAKVSATGDDAREGL 319
GYRB_STAAN      SGYATNLLTYANNIHTYEGGTHEDGFKRALTRVLNSYGLSSKIMKEEKDRLSGEDTREGM 330
                 *:   .:   ::*.*   :****   **: *:* .:* *   .     *. .    :*:* ***:

GYRB_MYCTU      AAVISVKVSEPQFEGQTKTKLGNTEVKSFVQKVCNEQLTHWFEANPTDAKVVVNKAVSSA 414
GYRB_ECOLI      IAVVSVKVPDPKFSSQTKDKLVSSEVKSAVEQQMNELLAEYLLENPTDAKIVVGKIIDAA 379
GYRB_STAAN      TAIISIKHGDPQFEGQTKTKLGNSEVRQVVDKLFSEHFERFLYENPQVARTVVEKGIMAA 390
                 *::*:*  :*:*..*** ** .:**:. *::   .* : .::   **   *: ** * : :*

GYRB_MYCTU      QARIAARKARELVRRKSATDIGGLPGKLADCRSTDPRKSELYVVEGDSAGGSAKSGRDSM 474
GYRB_ECOLI      RAREAARRAREMTRRKGALDLAGLPGKLADCQERDPALSELYLVEGDSAGGSAKQGRNRK 439
GYRB_STAAN      RARVAAKKAREVTRRKSALDVASLPGKLADCSSKSPEECEIFLVEGDSAGGSTKSGRDSR 450
                :** **::***:.***.* *:..******** . .*  .*:::*********:*.**:

GYRB_MYCTU      FQAILPLRGKIINVEKARIDRVLKNTEVQAIITALGTGIHD-EFDIGKLRYHKIVLMADA 533
GYRB_ECOLI      NQAILPLKGKILNVEKARFDKMLSSQEVATLITALGCGIGRDEYNPDKLRYHSIIIMTDA 499
GYRB_STAAN      TQAILPLRGKILNVEKARLDRILNNNEIRQMITAFGTGIGGD-FDLAKARYHKIVIMTDA 509
                 ******:***:******:*::*.. *:  :***:* **     ::  * ***.*::*:**

GYRB_MYCTU      DVDGQHISTLLLTLLFRFMRPLIENGHVFLAQPPLYKLKWQRSDPEFAYSDRERDGLLEA 593
GYRB_ECOLI      DVDGSHIRTLLLTFFYRQMPEIVERGHVYIAQPPLYKVKKGKQ-EQYIKDDEAMDQYQIS 558
GYRB_STAAN      DVDGAHIRTLLLTFFYRFMRPLIEAGYVYIAQPPLYKLTQGKQ-KYYVYNDRELDKLK-- 566
                **** ** *****:::* *  ::* *:*::*******:.  :.    :  .*.   *

GYRB_MYCTU      GLKAGKKINKEDGIQRYKGLGEMDAKELWETTMDPSVRVLRQVTLDDAAAADELFSILMG 653
GYRB_ECOLI      IA[169]SRRGLSIQRYKGLGEMNPEQLWETTMDPESRRMLRVTVKDAIAADQLFTTLMG 782
GYRB_STAAN      ---SELNPTPKWSIARYKGLGEMNADQLWETTMNPEHRALLQVKLEDAIEADQTFEMLMG 623
                            .* ********: .:******:*. * : :*.:.**   **: *  ***

GYRB_MYCTU      EDVDARRSFITRNAKDVRFLDV 675
GYRB_ECOLI      DAVEPRRAFIEENALKAANIDI 804
GYRB_STAAN      DVVENRRQFIEDNAVYA-NLDF 644
                : *: ** **  **   .  :*.
```

**Supplementary Figure S4. Sequence alignment of GyrB from *M. tuberculosis* (GYRB_MYCTU), *E. coli* and *S. aureus*.** Note the presence of an extra domain [169] within the *E. coli* Greek Key domain. The flexible linker region is ***underlined***. The residues in the Toprim domain are coloured red (see Figure 1). Sequences from UNIPROT.

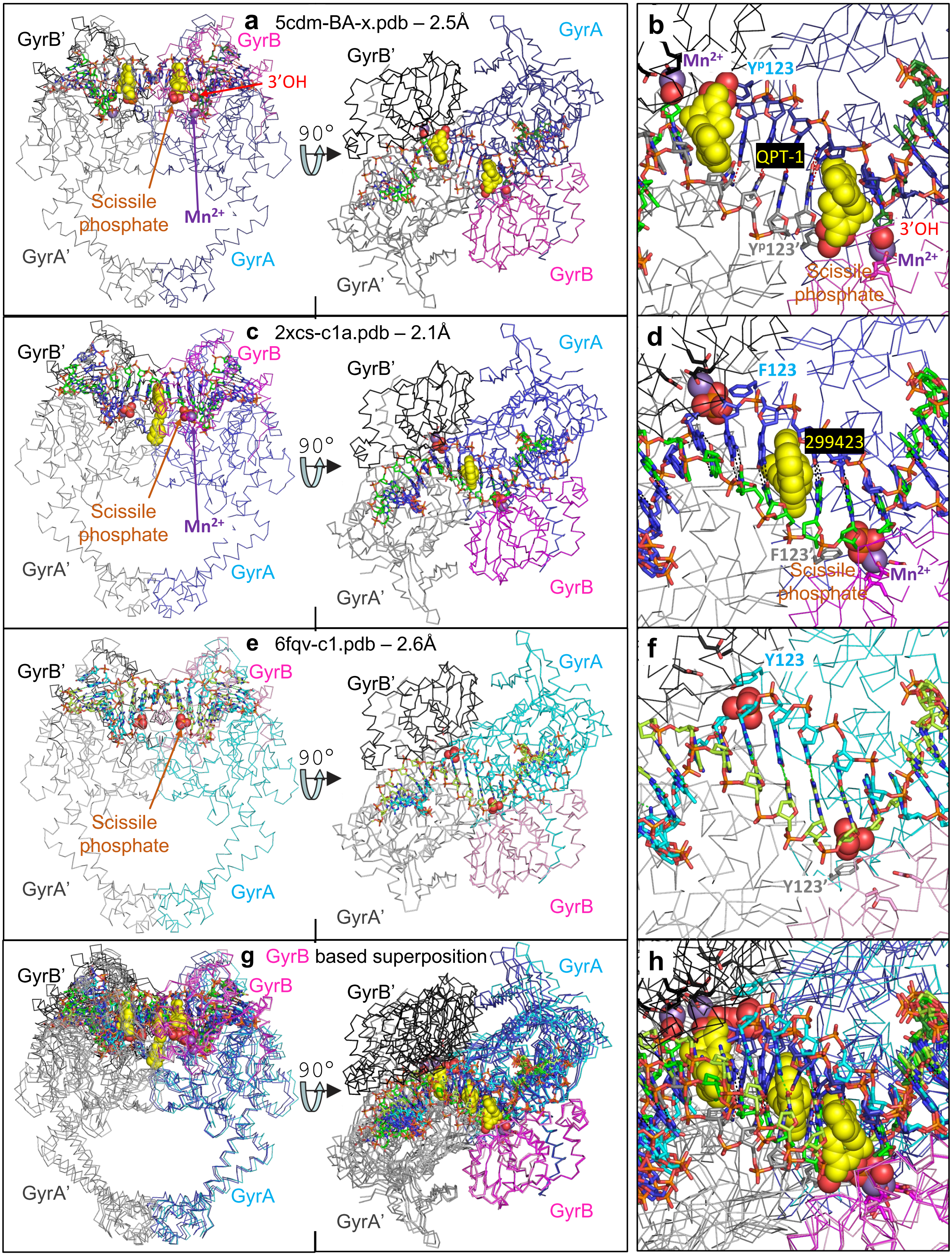


**Supplementary Figure S5. Catalytic $Mn^{2+}$ ions in crystal structures of *S. aureus* DNA gyrase. (a,b)** Views of a 2.5Å crystal structure of complex with QPT-1 and doubly cleaved DNA (pdb code: 5cdm). The catalytic metal is at the Y(B) site. Only one of the two scissile phosphates (shown as spheres) is labelled. **(c,d)** Views of a 2.1Å crystal structure of *S. aureus* DNA gyrase GyrB27–A56(GKdel) complex with GSK299423 and uncleaved DNA (pdb code: 2xcs). The catalytic metal is at the 3'(A) site contacting the scissile phosphate. The catalytic tyrosine has been mutated to a phenylalanine (Y123F). **(e,f)** Views of a 2.6Å crystal structure of a *S. aureus* DNA gyrase GyrB27–A56(GKdel) complex with uncleaved DNA. No metal ion was observed at any active site. **(g,h)** The three structures above are superposed based on the GyrB subunit. Coordinates available from the Research tab at: https://profiles.cardiff.ac.uk/staff/baxb.

```
                ---------- TOPRIM DOMAIN ---------
GYRA_MYCTU      MTDTTLPPDDSLDRIEPVDIEQEMQRSYIDYAMSVIVGRALPEVRDGLKPVHRRVLYAMF    60
GYRA_ECOLI      ------MSDL-AREITPVNIEEELKSSYLDYAMSVIVGRALPDVRDGLKPVHRRVLYAMN    53
GYRA_STAAN      ------MAELPQSRINERNITSEMRESFLDYAMSVIVARALPDVRDGLKPVHRRILYGLN    54
GYRA_BACSU      ------MSEQNTPQVREINISQEMRTSFLDYAMSVIVSRALPDVRDGLKPVHRRILYAMN    54
                        :    .:   :* .*:: *::********.****:***********:**.:

GYRA_MYCTU      DSGFRPDRSHAKSARSVAETMGNYHPHGDASIYDSLVRMAQPWSLRYPLVDGQGNFGSPG    120
GYRA_ECOLI      VLGNDWNKAYKKSARVVGDVIGKYHPHGDSAVYDTIVRMAQPFSLRYMLVDGQGNFGSID    113
GYRA_STAAN      EQGMTPDKSYKKSARIVGDVMGKYHPHGDSSIYEAMVRMAQDFSYRYPLVDGQGNFGSMD    114
GYRA_BACSU      DLGMTSDKPYKKSARIVGEVIGKYHPHGDSAVYESMVRMAQDFNYRYMLVDGHGNFGSVD    114
                  *   :: : **** *.:.:*:******:::*:::***** :. ** ****:***** .

GYRA_MYCTU      NDPPAAMRYTEARLTPLAMEMLREIDEETVDFIPNYDGRVQEPTVLPSRFPNLLANGSGG    180
GYRA_ECOLI      GDSAAAMRYTEIRLAKIAHELMADLEKETVDFVDNYDGTEKIPDVMPTKIPNLLVNGSSG    173
GYRA_STAAN      GDGAAAMRYTEARMTKITLELLRDINKDTIDFIDNYDGNEREPSVLPARFPNLLANGASG    174
GYRA_BACSU      GDSAAAMRYTEARMSKISMEILRDITKDTIDYQDNYDGSEREPVVMPSRFPNLLVNGAAG    174
                .*  ******* *:: :: *:: :: ::*:*:  ****  : * *:*::****.**:.*

GYRA_MYCTU      IAVGMATNIPPHNLRELADAVFWALENHDADEEETLAAVMGRVKGPDFPTAGLIVGSQGT    240
GYRA_ECOLI      IAVGMATNIPPHNLTEVINGCLAYIDDED----ISIEGLMEHIPGPDFPTAAIINGRRGI    229
GYRA_STAAN      IAVGMATNIPPHNLTELINGVLSLSKNPD----ISIAELMEDIEGPDFPTAGLILGKSGI    230
GYRA_BACSU      IAVGMATNIPPHQLGEIIDGVLAVSENPD----ITIPELMEVIPGPDFPTAGQILGRSGI    230
                ************:* *: :. :   .: *     ::  :*  : *******. * *  *

                              --------------- TOWER  DOMAIN ----------------
GYRA_MYCTU      ADAYKTGRGSIRMRGVVEVEE-DSRGRTSLVITELPYQVNHDNFITSIAEQVRDGKLAGI    299
GYRA_ECOLI      EEAYRTGRGKVYIRARAEVEVDAKTGRETIIVHEIPYQVNKARLIEKIAELVKEKRVEGI    289
GYRA_STAAN      RRAYETGRGSIQMRSRAVIEE-RGGGRQRIVVTEIPFQVNKARMIEKIAELVRDKKIDGI    289
GYRA_BACSU      RKAYESGRGSITIRAKAEIEQ-TSSGKERIIVTELPYQVNKAKLIEKIADLVRDKKIEGI    289
                  **.:***.: :*. . :*     *:  ::: *:*:***: .:* .**: *:: :: **

                ------------ TOWER  DOMAIN -----------
GYRA_MYCTU      SNIEDQSSDRVGLRIVIEIKRDAVAKVVINNLYKHTQLQTSFGANMLAIVDGVPRTLRLD    359
GYRA_ECOLI      SALRDES-DKDGMRIVIEVKRDAVGEVVLNNLYSQTQLQVSFGINMVALHHGQPKIMNLK    348
GYRA_STAAN      TDLRDETSLRTGVRVVIDVRKDANASVILNNLYKQTPLQTSFGVNMIALVNGRPKLINLK    349
GYRA_BACSU      TDLRDES-DRTGMRIVIEIRRDANANVILNNLYKQTALQTSFGINLLALVDGQPKVLTLK    348
                : :.*::  : *:*:**::::** ..*::****.:* **.*** *::*: .* *: : *.

                             --- COILED-COIL  DOMAIN ---   -- C-GATE DOMAIN -
GYRA_MYCTU      QLIRYYVDHQLDVIVRRTTYRLRKANERAHILRGLVKALDALDEVIALIRASETVDIARA    419
GYRA_ECOLI      DIIAAFVRHRREVVTRRTIFELRKARDRAHILEALAVALANIDPIIELIRHAPTPAEAKT    408
GYRA_STAAN      EALVHYLEHQKTVVRRRTQYNLRKAKDRAHILEGLRIALDHIDEIISTIRESDTDKVAME    409
GYRA_BACSU      QCLEHYLDHQKVVIRRRTAYELRKAEARAHILEGLRVALDHLDAVISLIRNSQTAEIART    408
                : :  :: *:  *: *** :.****. *****..*  **  :* :*  ** : *   *

                -------------------------------------- C-GATE DOMAIN --------
GYRA_MYCTU      GL----------------------------------IELLDIDEIQAQAILDMQLRRLAA    445
GYRA_ECOLI      ALVANPWQLGNVAAMLERAGDDAARPEWLEPEFGVRDGLYYLTEQQAQAILDLRLQKLTG    468
GYRA_STAAN      SL----------------------------------QQRFKLSEKQAQAILDMRLRRLTG    435
GYRA_BACSU      GL----------------------------------IEQFSLTEKQAQAILDMRLQRLTG    434
                .*                                       : * *******::*::*:.

                -- COILED-COIL  DOMAIN -
GYRA_MYCTU      LERQRIIDDLAKIEAEIADLEDILAKPERQRGIVRDELAEIVDRHGDDRRTRIIAA-DGD    504
GYRA_ECOLI      LEHEKLLDEYKELLDQIAELLRILGSADRLMEVIREELELVREQFGDKRRTEITAN-SAD    527
GYRA_STAAN      LERDKIEAEYNELLNYISELETILADEEVLLQLVRDELTEIRDRFGDDRRTEIQLGGFED    495
GYRA_BACSU      LEREKIEEEYQSLVKLIAELKDILANEYKVLEIIREELTEIKERFNDERRTEIVTSGLET    494
                **::::  :  .:   *::*  **..      ::*:**  : ::..*.***.*
```

**Supplementary Figure S6 part 1.  Sequence alignment of GyrA from *M. tuberculosis* (GYRB_MYCTU),  *E. coli, S. aureus and B. subtillis.*** Note the region in the C-gate of E.coli correlateds with the extra  domain within the *E.coli.* GyrB. The approximate positions of domains are indicated. Note many residues, including the catalytic **RY**, are in the winged helical domain (not shown). Sequences from UNIPROT.

```
                      ----------------- C-Domain ----------------------
GYRA_MYCTU  505 VSDEDLIAREDVVVTITETGYAKRTKTDLYRSQKRGGKGVQGAGLKQDDIVAHFFVCSTH  564
GYRA_ECOLI  528 INLEDLITQEDVVVTLSHQGYVKYQPLSEYEAQRRGGKGKSAARIKEEDFIDRLLVANTH  587
GYRA_STAAN  496 LEDEDLIPEEQIVITLSHNNYIKRLPVSTYRAQNRGGRGVQGMNTLEEDFVSQLVTLSTH  555
GYRA_BACSU  495 IEDEDLIERENIVVTLTHNGYVKRLPASTYRSQKRGGKGVQGMGTNEDDFVEHLISTSTH  554
                :. **** .*::*:*::. .* *    . *.:*.***:* ..    ::*:: ::.  .**

                ------------------------ C-Domain ------------------------
GYRA_MYCTU      DLILFFTTQGRVYRAKAYDLPEASRTARGQHVANLLAFQPEERIAQVIQIRGYTD-APYL  623
GYRA_ECOLI      DHILCFSSRGRVYSMKVYQLPEATRGARGRPIVNLLPLEQDERITAILPVTEFEE-GVKV  646
GYRA_STAAN      DHVLFFTNKGRVYKLKGYEVPELSRQSKGIPVVNAIELENDEVISTMIAVKDLESEDNFL  615
GYRA_BACSU      DTILFFSNKGKVYRAKGYEIPEYGRTAKGIPIINLLEVEKGEWINAIIPVTEFNA-ELYL  613
                * :* *:.:*:**  * *::**  * ::*  : * : .:  * *  :: :         :

                ------------------------ C-Domain ------------------------
GYRA_MYCTU      VLATRNGLVKKSKLTDFDSNRSGGIVAVNLRDNDELVGAVLCSAGDDLLLVSANGQSIRF  683
GYRA_ECOLI      FMATANGTVKKTVLTEFNRLRTAGKVAIKLVDGDELIGVDLTSGEDEVMLFSAEGKVVRF  706
GYRA_STAAN      VFATKRGVVKRSALSNFSRINRNGKIAISFREDDELIAVRLTSGQEDILIGTSHASLIRF  675
GYRA_BACSU      FFTTKHGVSKRTSLSQFANIRNNGLIALSLREDDELMGVRLTDGTKQIIIGTKNGLLIRF  673
                 .::* .*  *:: *::*    .  * :*:.: :.***:.. * .. .:::: : ..  :**

                ------------------------ C-Domain ------------------------
GYRA_MYCTU      SATDEALRPMGRATSGVQGMRFNIDDRLLSLNVVREG--TYLLVATSGGYAKRTAIEEYP  741
GYRA_ECOLI      KES--SVRAMGCNTTGVRGIRLGEGDKVVSLIVPRGD--GAILTATQNGYGKRTAVAEYP  762
GYRA_STAAN      PES--TLRPLGRTATGVKGITLREGDEVVGLDVAHANSVDEVLVVTENGYGKRTPVNDYR  733
GYRA_BACSU      PET--DVREMGRTAAGVKGITLTDDDVVVGMEILEEE--SHVLIVTEKGYGKRTPAEEYR  729
                  :   :* :*  ::**:*: :  .* ::.: : .      :* .*. **.***   :*

                ------------------------ C-Domain ------------------------
GYRA_MYCTU      VQGRGGKGVLTVMYDRRRGRLVGALIVDDDSELYAVTSGGGVIRTAARQVRKAGRQTKGV  801
GYRA_ECOLI      TKSRATKGVISIKVTERNGLVVGAVQVDDCDQIMMITDAGTLVRTRVSEISIVGRNTQGV  822
GYRA_STAAN      LSNRGGKGIKTATITERNGNVVCITTVTGEEDLMIVTNAGVIIRLDVADISQNGRAAQGV  793
GYRA_BACSU      TQSRGGKGLKTAKITENNGQLVAVKATKGEEDLMIITASGVLIRMDINDISITGRVTQGV  789
                 ..*. **: :    ...* :*    . . .::  :* .* ::*    ::   ** ::**

                ---- C-Domain -----
GYRA_MYCTU      RLMNLGEGDTLLAIARNAEESGDDNAVDA---NGADQTGN--------------------  838
GYRA_ECOLI      ILIRTAEDENVVGLQRVAEPVDEEDLDTI---DGSAAEGDDEIAPEVDVDDEPEEE----  875
GYRA_STAAN      RLIRLGDDQFVSTVAKVKEDAEDETNEDEQSTSTVSEDGTEQQREAVVNDETPGNAIHTE  853
GYRA_BACSU      RLIRMAEEEHVATVALVEKNEEDENEEEQEEV----------------------------  821
                 *:. .: : :  :    :   ::

GYRA_MYCTU      ------------------------------------  838
GYRA_ECOLI      ------------------------------------  875
GYRA_STAAN      VIDSEENDEDGRIEVRQDFMDRVEEDIQQSSDEDEE  889
GYRA_BACSU      ------------------------------------  821
```

**Supplementary Figure S6 part 2. Sequence alignment of GyrA from *M. tuberculosis* (GYRB_MYCTU), *E. coli, S. aureus and B. subtillis* – continued.** Note this is a continuation of the alignment shown in Figure S5 part 1. This alignment in this figure (part 2) shows the short linker region (<10 amino acids), the C-terminal DNA-wrapping domain, and additional C-terminal amino acids (variable length). The approximate extent of the C-terminal wrapping domain in *E. coli* is indicated above the alignment. Aligments from Custalw.

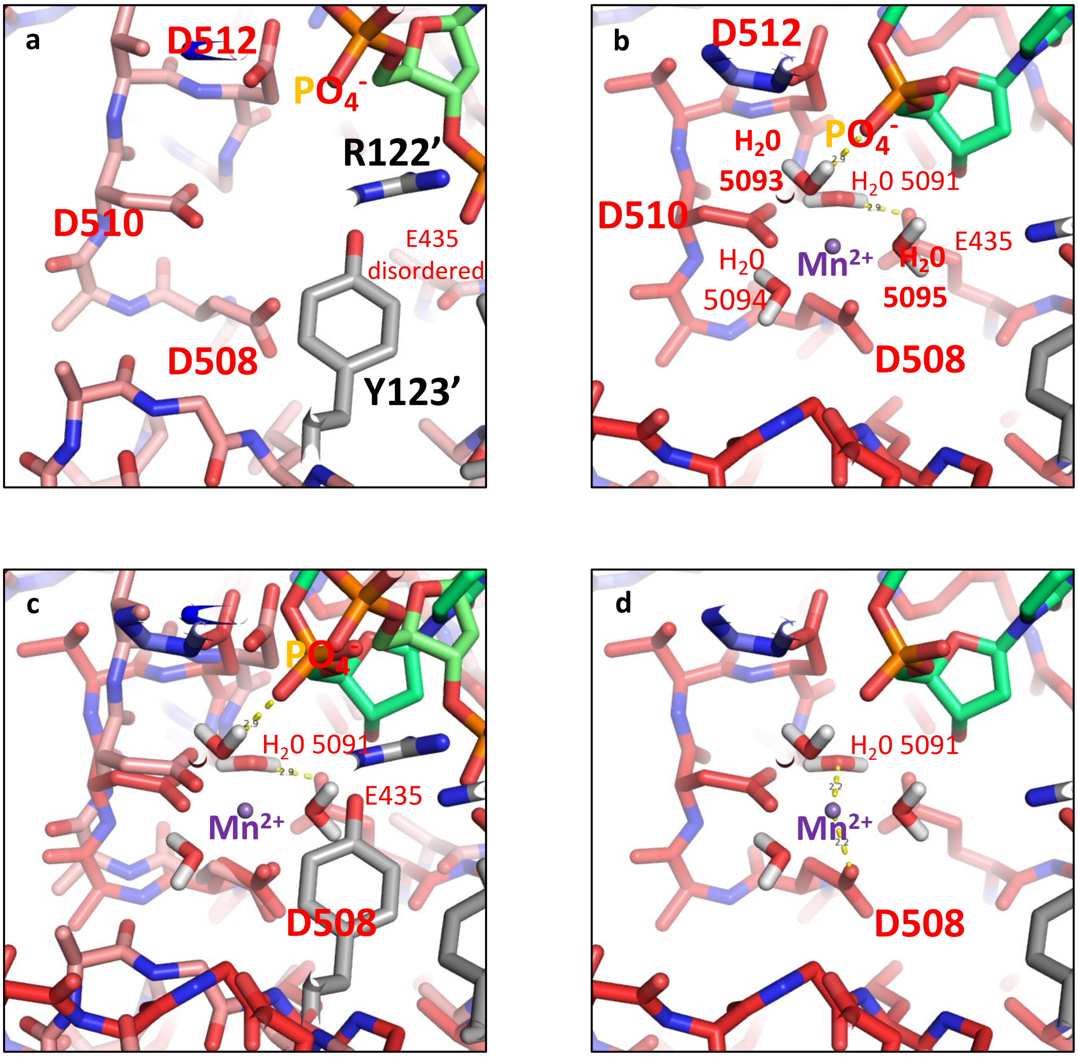


**Supplementary Figure S7. The presence of an $Mn^{2+}$ ion at the Y(B) site seems to order Glu B435 and attract the phosphate before the scissile phosphate towards water 5093.**
**(a)** A view of the 2.6Å crystal structure of a *S. aureus* DNA gyrase GyrB27–A56(GKdel) complex with uncleaved DNA (6fqv-c1.pdb). No metal ion is observed. Note the side-chain of Glu B435 is disordered. (**b**) A view of the Y(B) site metal in the 2.5Å structure of a complex with QPT-1 (the progenitor of zoliflodacin) and doubly cleaved DNA (pdb code: 5cdm). Hydrogen bonds are shown (dashed yelow lines) between water 5093 and the phosphate ($PO_4^-$) before the cleavage site and between water 5091 and the side-chain of Glu B435. Hydrogens have been modelled (Maestro) on the four waters coordinating the $Mn^{2+}$ ion. (**c**) The structures in panels a and b are shown superposed (based on GyrB). Note the ordering of Glu B435 and the relative movement of the labelled phosphate. (**d**) A view of the Y(B) site metal in the 2.5Å structure of a complex with QPT-1 (the progenitor of zoliflodacin) and doubly cleaved DNA (pdb code: 5cdm). Interactions are shown (dashed yellow lines each of length 2.2Å) between water 5091 and the $Mn^{2+}$ ion and between the side-chain of Asp B508 and the $Mn^{2+}$ ion.

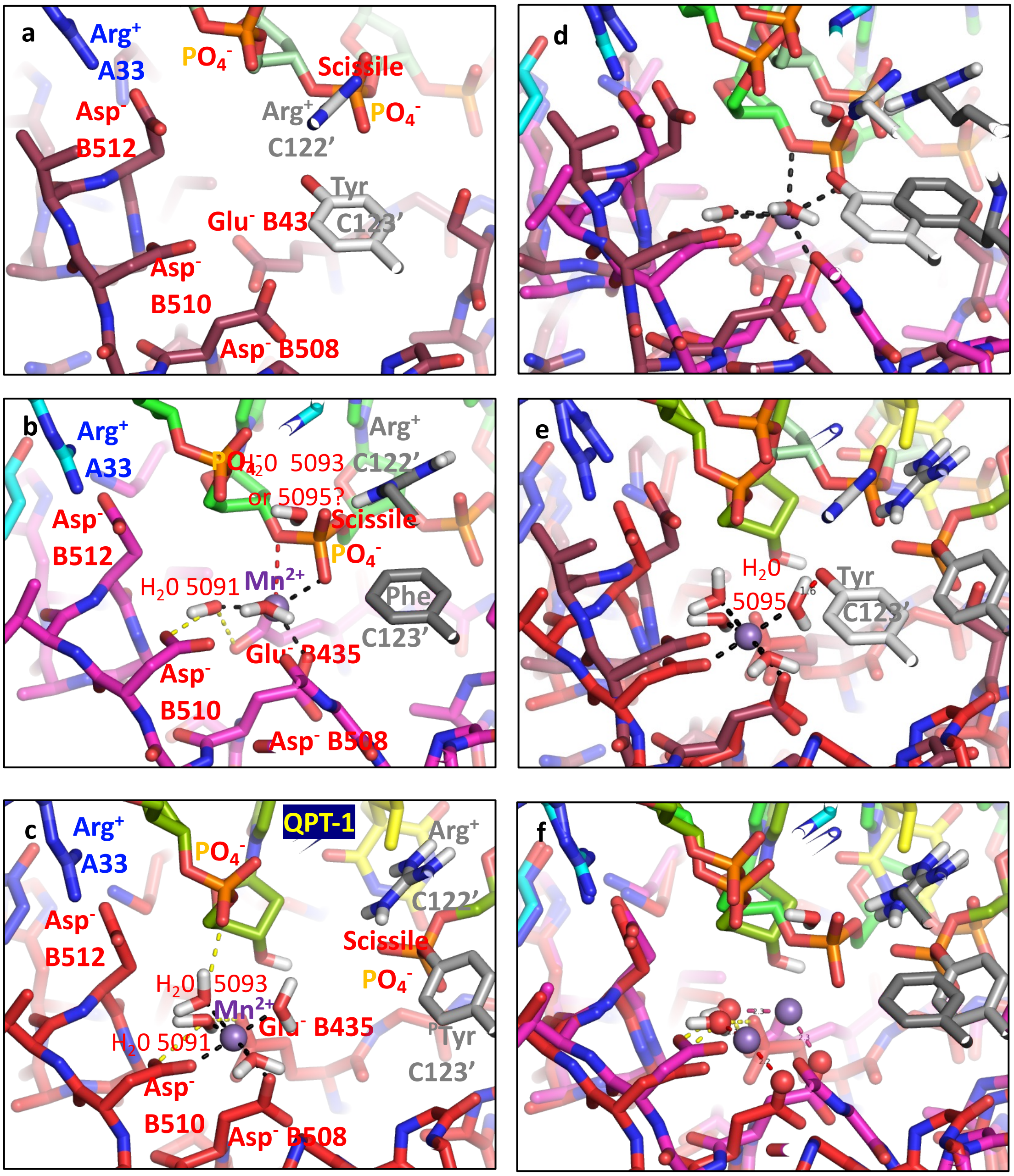


**Supplementary Figure S8. A comparison of 6fqv-c1-one-DNA.pdb with 2xcs-v2-two-DNA.pdb and 5cdm-v2-two-DNA.pdb.**

**(a)** A view of **6fqv-c1-one-DNA.pdb** . Note the side-chain of Glu B435 is disordered in the crystal structure (6fqv-c1.pdb) and is modelled in (see Supplementary Figure S6). (**b**) A similar view of **2xcs-v2-two-DNA.pdb** which has a single metal ion at the 3’(A) site. Hydrogens have been modelled (Maestro) onto the two waters coordinating the catalytic metal, and a third water within H-bonding distance of the scissile phosphate. Hydrogens (white atoms) are also shown on Arg C122’. (**c**) A similar view of **5cdm-v2-two-DNA.pdb**. For clarity hydrogens are only shown on the four waters coordinating the metal ion ($Mn^{2+}$), the 3’ OH, and Arg C122’. (**d**) Structures from panels a and b are shown superposed. Note how the two $PO_4^-$ labelled groups from the DNA backbone appear to have moved closer to the metal ion in **2xcs-v2-two-DNA.pdb** cf **6fqv-c1-one-DNA.pdb**. Note also the end of Arg C122’ from **6fqv-c1-one-DNA.pdb** appears to occupy a similar position to that of the water near the scissile phosphate in **2xcs-v2-two-DNA.pdb**. (**e**) Structures from panels a and c are shown superposed. Note the terminal oxygen of the cataytic tyrosine, Tyr C123’ from **6fqv-c1-one-DNA.pdb** is within 1.6Å of the oxygen in water 5095 in **5cdm-v2-two-DNA.pdb** (**f**) Structures from panels b and c are shown superposed. Note three ‘heavy atoms’ the oxygen from water 5091, the catalytic metal ($Mn^{2+}$) ion and the OD2 oxygen from Asp 508 are shown as spheres in both structures. Other waters coordinating the catalytic metal ($Mn^{2+}$) ion are not shown for clarity.

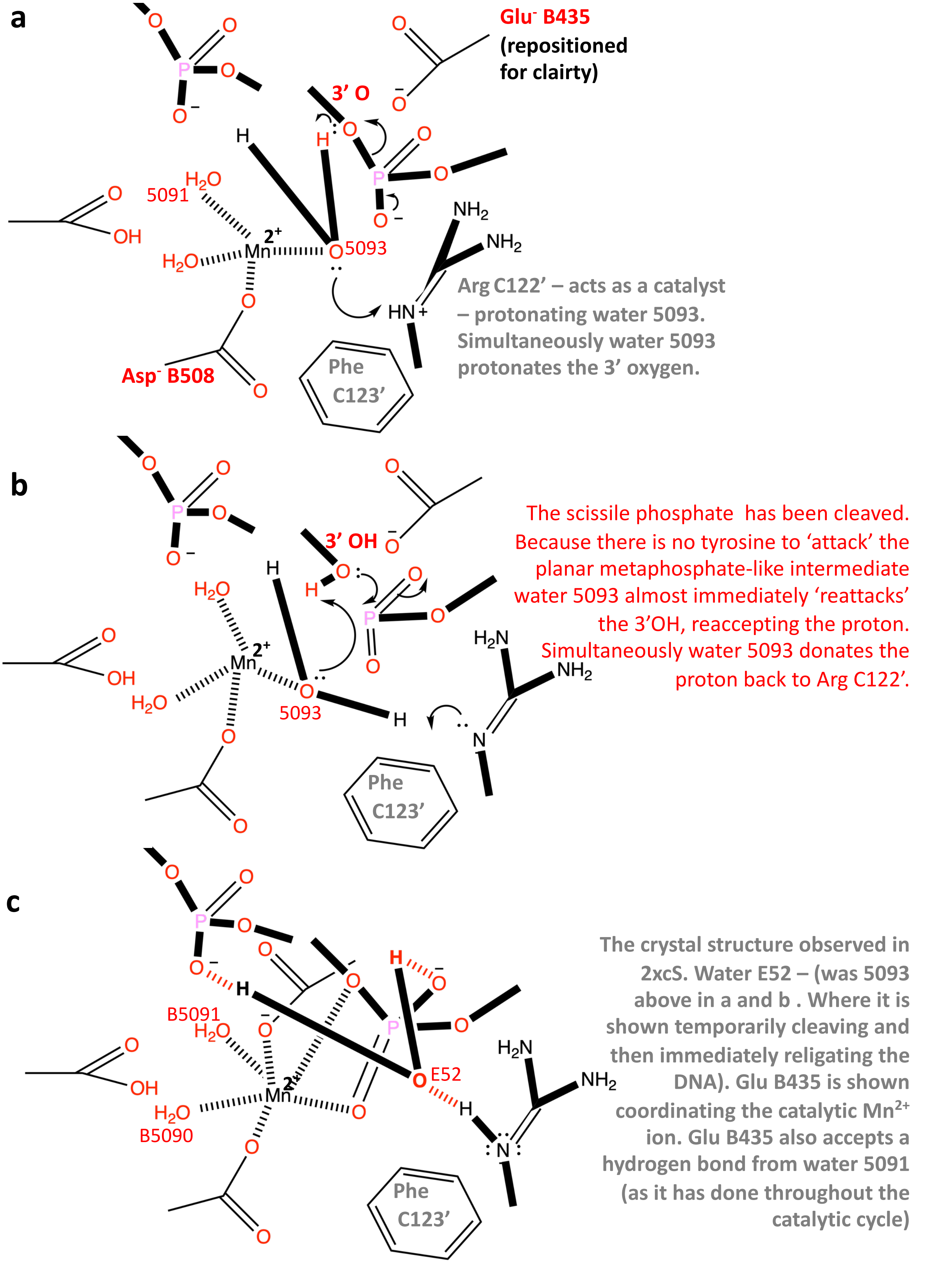


**Supplementary Figure S9. Could water 5093 have momentarily cleaved the DNA?**
**(a) W**ater 5093 is shown poised for protonation by the catalytic arginine (compare with figure 5 panel b). (**b**) Water 5093 has momentarily cleaved the DNA. Because the catalytic tyrosine has been mutated to a phenylanine it cannot 'attack' the highly reactive metaphosphate-like intermediate. So water 5093 reaccepts the hydrogen from the 3'OH and donates a hydrogen back to Arg C122'. (**c**) The crystal structure of 2xcs is drawn showing hydrogen bonds to water E52 as red hashed lines and the distorted octahedral coordination sphere of the Mn2+ ion as hashed black lines.
